# Pre-synaptic facilitation of corticospinal output improves dexterity and fine motor control after stroke

**DOI:** 10.64898/2026.11.06.764958

**Authors:** Arianna Damiani, Nicolo’ Macellari, Zikai Xu, Lucy Liang, Catherine Jerzerc, Lilly W. Tang, Katie Jones, Miles Uribe, Chaitanya Goswami, Kelly A. Hoffmann, Tyler Simpson, Erinn M. Grigsby, Isabella Naddeo, Peter Gerszten, Marco Capogrosso, Jorge Gonzalez-Martinez, Elvira Pirondini

## Abstract

The corticospinal tract (CST) is a neural pathway that supports skilled use of the arm and hand. Lesions of corticospinal axons result in muscle weakness, deficits of gross motor function, and loss of proximal and distal dexterous control for which effective treatments are still lacking. Here we exploited excitatory connections from motor thalamus to primary motor cortex to potentiate surviving descending signals and restore volitional arm and hand control in three monkeys with corticospinal tract lesion, using deep brain stimulation. Stimulation targeted to thalamocortical projections facilitated upper-limb movements, enabling monkeys with arm and hand paralysis to perform unconstrained 3D reach and fine dexterous tasks. Stimulation improved strength, hand gross motor function, and digits control. Electrophysiology suggested that effects were mediated by and more effective when targeting fibers with monosynaptic connection to the corticospinal motoneurons. Stimulation was delivered in continuous with FDA-clinically approved parameters holding realistic promises of clinical translation.

## Introduction

Stroke lesions reduce the cortical descending drive that reaches spinal motoneurons, causing the stereotypical manifestation of hemiparesis characterized by loss of strength (that is, loss of the ability to activate muscles)^1,2^; loss of proximal and distal dexterity (that is, loss of the ability to flexibly and independently control muscles and joints)^3–5;^ and loss of hand gross motor function^6,7^. Nowadays, there are more than 6 million people in the USA also suffering from such conditions^8,9^.

While high-dose motor rehabilitation can improve these symptoms^10,11^, the dose delivered in clinical practice is approximately ten times lower^4,12^, resulting in limited efficacy, particularly for the hand^13,14^. For this reason, in the past, several neurostimulation approaches have been proposed to catalyze the effects of the training. These include targeting the cortex directly (using non-invasive techniques or surgically implanted electrode grids) or indirectly, via stimulation of the cerebello-cortical pathway^15–18^. In both cases, the stimulation is applied before or throughout the training to increase ipsilesional cortical excitability and promote neuroplasticity. Similarly, vagus nerve stimulation has been shown to enhance the reorganization potential of the brain via cholinergic and monoaminergic modulation of motor cortex neurons, when stimulation is paired with repetitive movements^19,20^. While these technologies could improve strength, arm dexterity, and gross motor function over time, leading, in some cases, to clinical meaningful changes, effects on distal dexterity and digits control remain rather limited. Compared to the arm, these functions are mainly controlled by the corticospinal tract (CST)^21–23^, i.e., the monosynaptic pathway between layer V corticospinal neurons in the motor cortex and the spinal motoneurons, with little redundancy from other descending pathways. Yet, previous approaches were designed to enhance cortical reorganization rather than to increase activation of this pathway. If fine motor control requires a large corticospinal drive during movement execution, this mismatch could explain their limited effects on distal function. This raises the question of whether strong and continuous excitatory inputs to corticospinal neurons can potentiate descending drive to spinal motoneurons (i.e., CST output) improving hand function after stroke.

The motor thalamus (mTh) provides monosynaptic excitatory inputs to layer V corticospinal neurons in the primary motor cortex (M1) and therefore represents a candidate target for this. We previously demonstrated in anesthetized monkeys that stimulation of this thalamic nucleus could increase CST output^24^. Specifically, we showed that motor evoked potentials (MEPs) elicited by stimulation of the CST were larger when the mTh was stimulated concurrently at low-frequency (50-80Hz). The increase in MEPs was immediate, i.e., it appeared immediately when the stimulation was turned ON, contrary to what reported in other deep brain stimulation (DBS) approaches in which the potentiation was present only when the stimulation was priorly ON for at least 45 minutes^18^. Importantly, when delivered alone (ie., not paired with CST stimulation), mTh stimulation was not generating artificial motor output.

Such immediate effects and the absence of artificial response could define a novel mode of intervention in which the individual retains volitional control but the stimulation amplifies the output (assistive effects). This represents a switch from the standard approaches designed to induce only lasting therapeutic effects, i.e., improvements that persist over time in the absence of stimulation, and from the idea of replacement effects, i.e., the stimulation directly generates an overt motor response. In this latter case, the stimulation overwrites the natural cortical activity^25,26^ and therefore requires a complex technology to detect motor intention and to deliver spatio-temporal stimulation patterns to activate muscles at the correct moment in time^27–29^. Such solutions are difficult to implement with clinically available stimulators and have therefore rarely moved beyond small laboratory studies.

Here we hypothesized that continuous stimulation of the motor thalamus would increase CST output in animals with chronic CST lesions, and that this would immediately improve upper-limb movements. We first demonstrated that the increased CST output was driven by thalamocortical monosynaptic excitatory inputs to corticospinal neurons. Indeed, the effects were stronger when stimulating the mTh than the nuclei projecting to it, and disappeared when dissecting the motor cortex. We then developed a controlled lesion of the posterior limb of the internal capsule in n=3 monkeys, which selectively interrupted corticospinal axons, and verified that the absence of these fibers led to chronic muscle weakness, loss of gross hand motor function and loss of finger control. mTh DBS immediately enabled reaching and grasping movements that were otherwise impossible when impairments were most severe, and later improved end-effector stabilization, grip control and strength, and dexterous hand movements. Finally, we showed that such improvements could be obtained with stimulation configurations already available in the clinic, without the need for detection of motor intention or phase-dependent stimulation patterns as in previous works, facilitating translation to humans.

## Results

### Hypothesis: motor thalamus stimulation potentiates CST output leading to improved volitional control

We previously demonstrated in anesthetized monkeys that continuous stimulation of the ventral laterolateral motor thalamus (corresponding to the ventral intermediate (VIM)/ventral oral posterior (VOP) target used clinically^30^ and that we will here refer to as mTh for the rest of the manuscript) at low-frequency (50–80 Hz, i.e., below the >130 Hz typically used in clinical DBS) potentiates corticospinal output both in intact animals and in animals with acute lesions^24^. Specifically, the peak-to-peak amplitude of motor evoked potentials (MEPs) elicited by stimulation of the CST was larger when the mTh was stimulated concurrently at low-frequency. We now hypothesize that this potentiation would lead to functional improvements of volitional control after CST lesions particularly for fine motor control (**Figure 1a**).

**Figure 1:**
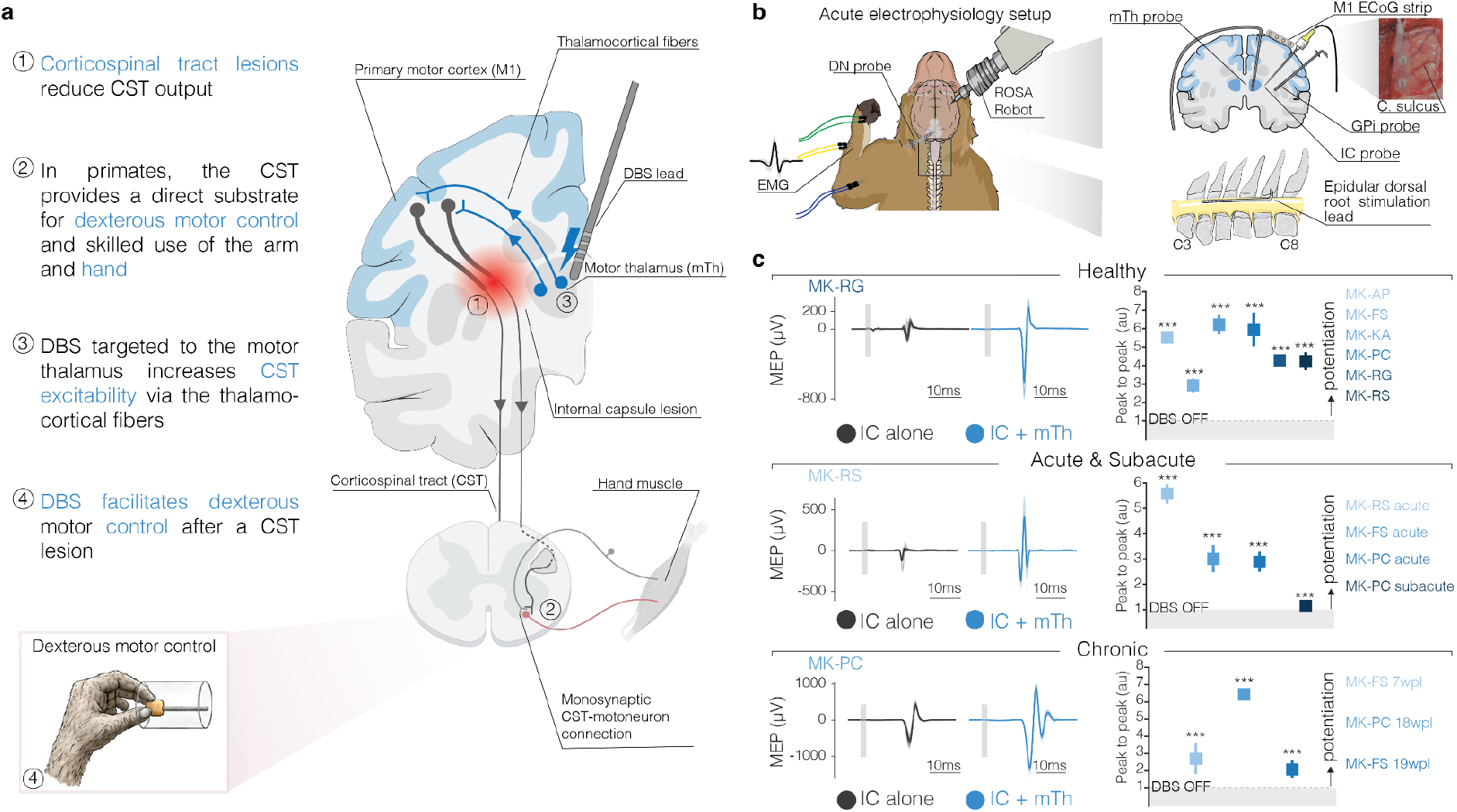
mTh stimulation at low-frequency potentiate corticospinal output. (**a**) Study hypothesis. (**b**) Experimental setup for anesthetized experiments. Animals were implanted with a stimulation probe in the internal capsule (IC) and a stimulation probe in the mTh, GPi and DN using the ROSA robot and a subdural strip over M1. An epidural lead was implanted at the C4-C7 spinal segment to elicit H-reflexes and EMG needle electrodes were inserted in arm, hand, finger muscles. (**c**) <u>Left:</u> Example of motor evoked potential (MEP) in a finger muscle with IC stimulation alone (black) and IC stimulation paired with mTh stimulation at low-frequency (50-80 Hz, blue). *<u>Right:</u>*Normalized peak to peak MEP amplitude change (IC + low-frequency mTh stimulation relative to IC alone) of one finger or hand muscle for each animal at different stages of stroke recovery (healthy: for MK-FS APB, for all other monkeys FDM; acute and subacute: for MK-RS and MK-PC acute FDM, for MK-FS and MK-PC subacute APB; chronic: ECR for MK-FS 7wpl, FCR for MK-PC 18wpl and MK-FS 19 wpl). Data shown as mean ± 95% CI. Results for all muscles of all monkeys in **Extended Data Figure 1**, and stimulation parameters in **Supplementary Table 2**. For all panels, statistical significance was assessed with two-tail bootstrapping with Bonferroni correction: p<0.05 (*), p<0.01 (**), p<0.001(***).

Before testing this hypothesis in behaving animals, we performed electrophysiology experiments to support our mechanism of action (n=7 monkeys, **Supplementary Table 1**). First, we verified that potentiation of corticospinal output was still present after subacute and chronic CST lesions. For this, we paired low-frequency (50-80Hz) stimulation of the motor thalamus with stimulation of the CST (either via the internal capsule or the motor cortex). Leveraging and expanding our previously described surgical robotic method^31^ (ROSA ONE® Robot Assistance Platform, ROSA robot, Zimmer Biomet, Warsaw, Indiana, USA) and experimental setup^24^ (**Figure 1b**), we implanted a stimulation probe in the mTh (**Extended Data Figure 1a**). High-resolution post-mortem structural MRI confirmed the accurate location of the electrode within the mTh (average distance between the implantation location and the mTh nucleus over animals: 1.49 mm ± 0.28 mm (mean±SE), in line with human studies^32^. A second probe was implanted in the posterior limb of the internal capsule (IC) where the arm and hand CST fibres are located and a 6 contact ECoG strip over the hand knob of M1 (**Figure 1b**). Using the IC probe, we performed a controlled lesion of the corticospinal axons by thermocoagulating them (85 °C for 60 seconds)^24,33^, which allowed us to test potentiation in intact animals (n=7) and immediately after the lesion (n=3 acute). Potentiation at later timepoints was assessed in animals that had undergone the same lesion as part of the behavioural protocol described below (n=1 subacute, 3 weeks post-lesion (WPL); n=3 chronic: 7, 18 and 19 WPL). Specifically, in each animal, we stimulated the IC at 1-2 Hz to elicit short-latency MEPs (10-1ms) in arm, hand, and finger muscles, consistent with the activation of corticospinal axons with direct connections to spinal motoneurons. We then paired stimulation of the IC with continuous mTh stimulation at low-frequency (50-80 Hz) (**Supplementary Table 2** for all stimulation parameters). As we previously had shown^24^, the peak-to-peak amplitudes of MEPs were immediately and significantly increased (up to ∼6 folds) when pairing single pulse stimulation of the IC with continuous stimulation of mTh in all tested animals and muscles (**Figure 1c, Extended Data Figure 1b**). Importantly, potentiation was present also weeks after the lesion. Crucially, we also confirmed that MEPs generated by direct activation of the motor cortex were potentiated by mTh stimulation (**Extended Data Figure 2a**). For this, we stimulated (burst of 10 pulses at 300-400 Hz, with inter-burst frequency of 2 Hz) from the ECoG contact placed over the hand knob of M1 to generate MEPs in hand and fingers muscles. We then paired cortical stimulation with continuous low-frequency (50-80 Hz) mTh stimulation. As with IC stimulation, MEPs amplitude was significantly higher when thalamic stimulation was turned ON (MK-RG hand +156.4%, finger +40.7%; MK-AP arm +127.4%, hand +209.5%, finger +116.4%, **Extended Data Figure 2b**).

This demonstrated that low-frequency mTh stimulation increased corticospinal output. Importantly, the potentiation was also present from weeks to months after the lesion, suggesting that mTh stimulation could be used to improve volitional control in animals with acute and chronic injuries.

### Specificity of the stimulation

The mTh receives inputs from the basal ganglia via the globus pallidus internus (GPi) and from the cerebellum through the dentate nucleus (DN)^34^ (**Figure 2a**). Previous works showed that stimulation of these two nuclei also potentiates upper-limb MEPs^35,36^. However, these nuclei connect to the layer V corticospinal neurons disynaptically via the pallido-thalamo-cortical and the cerebello-thalamo-cortical circuit. We, therefore, questioned which of these three nuclei would produce the strongest MEP potentiation, hypothesizing that the effects would be larger for the mTh, which is only one synapse away from the corticospinal neurons.

**Figure 2:**
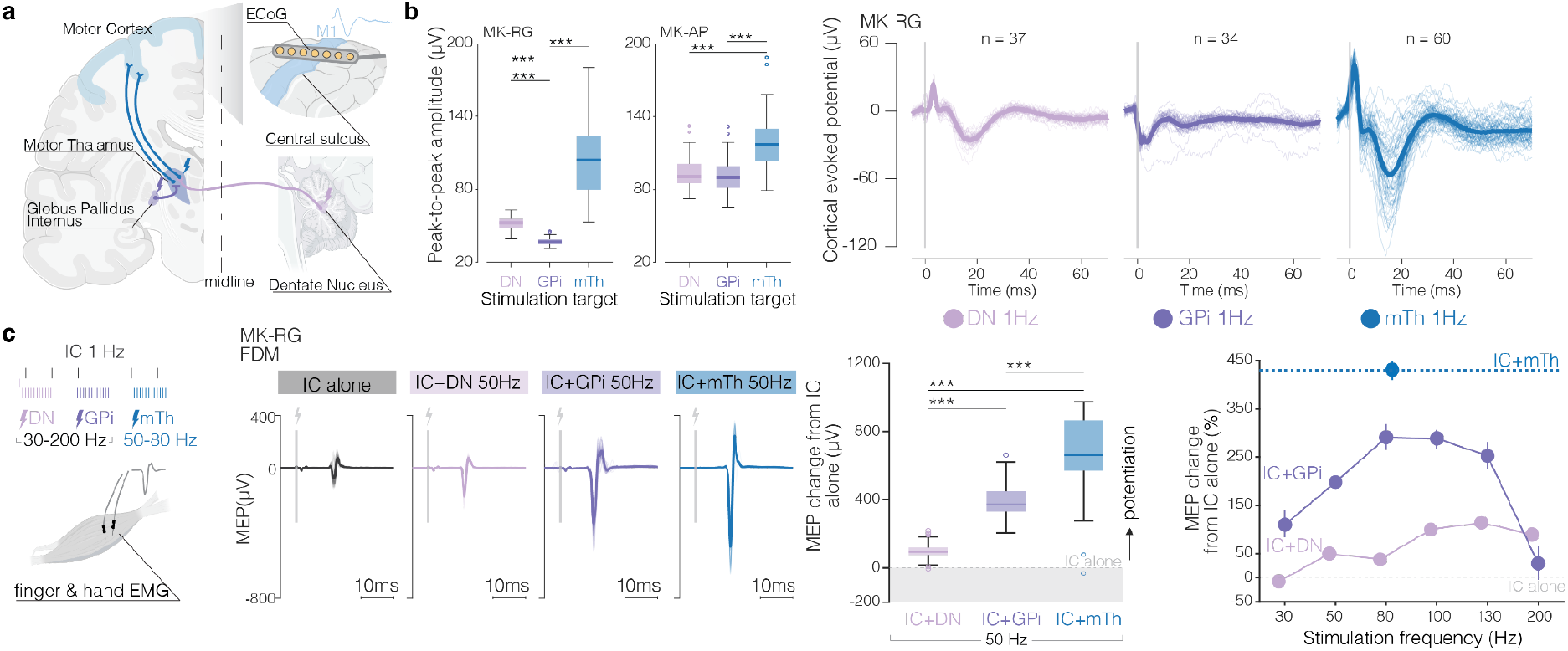
mTh stimulation maximally potentiates motor output compared to disynaptic targets. (**a**) Schematic of anatomical pathways and experimental setup used to test the specificity of mTh (blue) stimulation compared to DN (pink) and GPi (purple). On the right, a schematic of ECoG strip placed over M1 to record cortical evoked potentials. (**b**) <u>Left</u>: Box-plots of peak-to-peak amplitudes of cortical evoked potentials in M1 elicited by stimulation of the DN, GPi or mTh individually. <u>Right</u>: Representative cortical evoked responses for MK-RG. (**c**) <u>Left</u>: experimental protocol to compare the potentiation of motor output induced by DN, GPi and mTh stimulation. Stimulation of these nuclei was paired with 1 Hz stimulation of the IC, and MEPs were analyzed. <u>Middle</u>: example traces (MK-RG, FDM muscle) of MEPs for IC stimulation alone, IC+DN stimulation at 50Hz, IC+GPi stimulation at 50Hz, IC+mTh stimulation at 50Hz. For these traces, box plot of MEP peak-to-peak amplitude change with and without stimulation of the different nuclei, relative to IC alone baseline (median subtracted, y=0 line = IC alone). <u>Right</u>: percentage change in MEP amplitude relative to IC alone as a function of stimulation frequency for stimulation of the DN (pink) and the GPi (purple) and compared to the maximum potentiation achieved with low-frequency mTh stimulation (dashed blue line). Data shown as mean ± 95% CI. For all boxplots, the whiskers extend to the maximum spread not considering outliers, central, top, and bottom lines represent median, 25th, and 75th percentile, respectively. For all panels, statistical significance was assessed with two-tail bootstrapping with Bonferroni correction: p<0.05 (*), p<0.01 (**), p<0.001(***). Data from all muscles and all animals shown in **Extended Data Figure 3**, stimulation parameters in **Supplementary Table 2**.

For this, we leveraged our high-throughput electrophysiology set-up to implant one additional stimulating probe in the GPi and one in the DN (GPi: n=2 animals; DN: n=3 animals, average distance between the implantation location and each nucleus over animals: 0.90 mm ± 0.01 mm (mean±SE) for DN, 0.70 mm ± 0.05 mm (mean±SE) for GPi, **Extended Data Figure 3a**). We first stimulated each nucleus individually (1 Hz, 2-2.5 mA) while recording cortical evoked potentials (CEPs) from the ECoG strip over M1. Although stimulation of all nuclei resulted in reliable CEPs, CEP amplitude was significantly larger with mTh stimulation (**Figure 2b**, median CEP amplitude MK-RG: DN 52.33, GPi 36.39, mTh 104.1 uV; MK-AP: DN 90.69, GPi 90.07, mTh 116.7 uV; p<0.001), likely because of the monosynaptic connection between this thalamic area and the motor cortex. We then paired stimulation of each nucleus at 50 Hz with IC stimulation at 1 Hz, and recorded upper-limb MEPs. Stimulation amplitudes of the nuclei were chosen as the value in which MEPs peak-to-peak reached a plateau, and no direct CST activation or change in vital parameters was reported (see Methods, **Supplementary Table 2**, **Extended Data Figure 3b**). While all targets led to a sustained potentiation of the MEPs, peak-to-peak amplitudes were significantly larger when stimulating the mTh (**Figure 2c**). Because previous studies reported an optimal stimulation frequency of around 30 Hz for GPi and DN^18^, we compared the potentiation obtained with GPi and DN stimulation at different frequencies (30-100 Hz) to that obtained with mTh stimulation at 50-80 Hz, i.e., the frequency range we previously identified as optimal^24^. MEPs potentiation obtained with mTh stimulation was significantly stronger than the maximum potentiation obtained with DN or GPi stimulation at any tested frequency (for each nucleus maximum percentage change stimulation ON/OFF, MK-RG DN: 113.47%, GPi: 290%, mTh: 430.45%; MK-AP DN: 45.40%, GPi: 267.54%, mTh: 786.59%; MK-MA DN: 1371.16%, mTh: 1843.13%, **Figure 2c**, **Extended Data Figure 3c**), indicating that the stronger effects of mTh stimulation was not due to a suboptimal choice of stimulation parameters for the other nuclei.

Overall, these results confirm the mTh as the optimal stimulation location. We attribute the stronger MEP potentiation to the monosynaptic connections between the mTh and M1, whereas GPi and DN reach M1 only disynaptically, through the thalamus.

### Thalamocortical synaptic inputs mediate the potentiation of corticospinal output

We then sought to demonstrate that, indeed, the potentiation of corticospinal output was mediated by excitatory synaptic inputs from thalamocortical afferents onto the corticospinal neurons (H1). Alternatively, mTh stimulation could potentiate MEPs through off-target effects: direct activation of CST axons passing near the mTh, which would directly increase excitability of spinal motoneurons (H2); or spinal excitation arising from other poly-synaptic descending pathways or from subthreshold neuromodulation factors (H3). To distinguish between these alternative hypotheses (H1, H2 and H3), we designed a series of electrophysiology experiments (**Figure 3a**).

**Figure 3:**
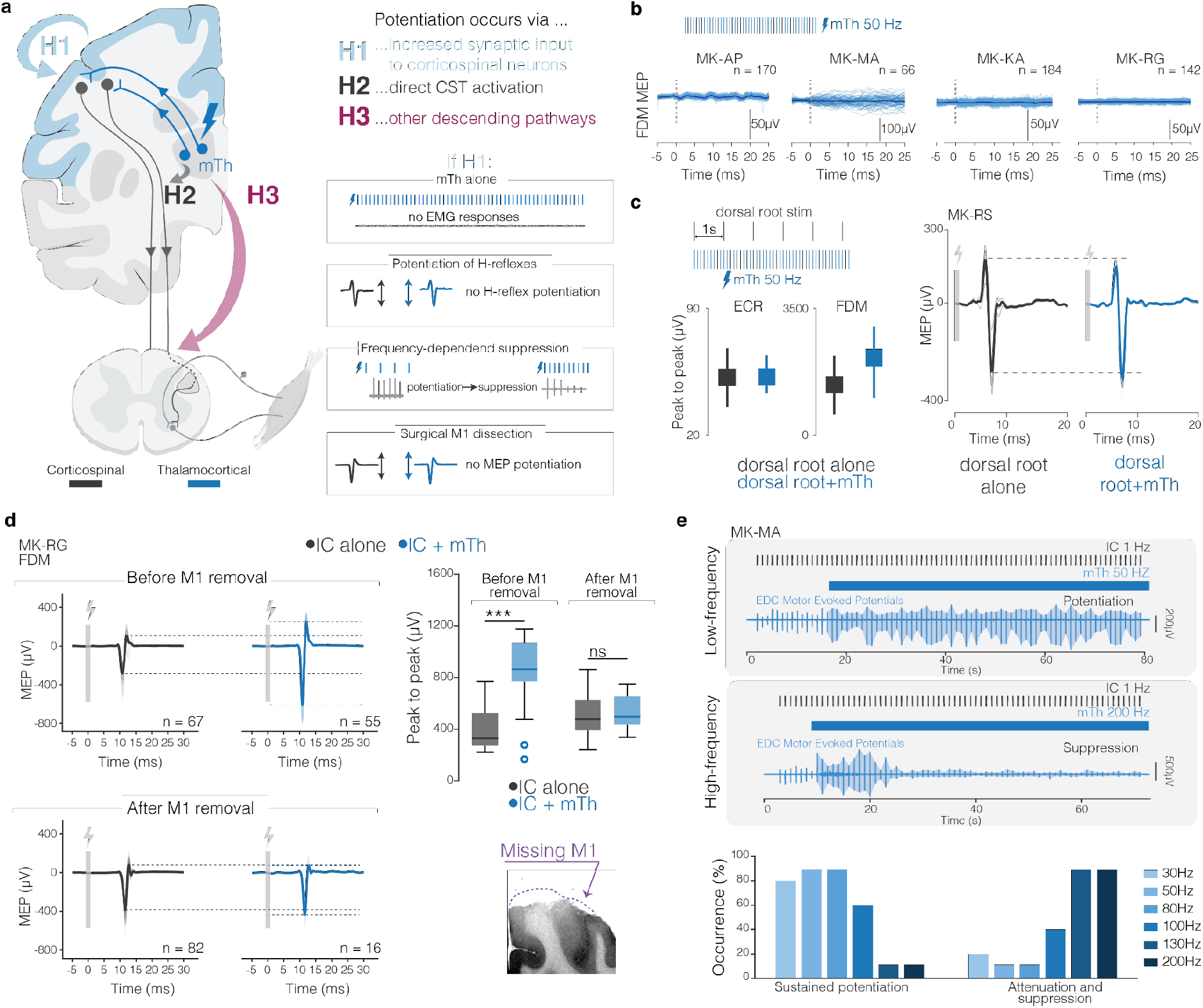
MEP potentiation results from increased M1 excitability, mediated by excitatory synaptic inputs from thalamocortical afferents. **(a)** Potential hypothesis that could explain the observed increase in MEPs with mTh stimulation. (**b**) Examples of one muscle (FDM) for all animals with mTh stimulation alone at 50 Hz. No MEPs were elicited. (**c**) <u>Top left</u>: stimulation protocol to test modulations in spinal excitability. We sent pulses of dorsal root stimulation to elicit H-reflex, and then paired it with low-frequency mTh stimulation. <u>Bottom Left</u>: peak-to-peak amplitude of H-reflex without (grey) and with (blue) mTh stimulation for example muscles of hand (ECR) and finger (FDM). Data shown as mean ± 95% CI. <u>Right:</u> example of H-reflex for dorsal root stimulation alone (grey) and dorsal root stimulation+mTh stimulation (blue) (MK-RS, FDM). **(d)** <u>Left</u>: example of MEP traces for IC alone (grey) and IC+mTh stimulation (blue) before and after disconnection of M1 (top and bottom panel, respectively). <u>Right</u>: Boxplot of peak-to-peak amplitudes of MEPs. For all boxplots, the whiskers extend to the maximum spread not considering outliers, central, top, and bottom lines represent median, 25th, and 75th percentile, respectively. On the bottom, an example of post-mortem MRI showing dissected M1. (**e**) <u>Top</u>: example of frequency-dependent MEPs modulation. In the top panel, IC stimulation (black pulses) is paired with 50 Hz mTh stimulation and representative MEP recording shows stable potentiation over time. On the contrary, in the bottom panel mTh stimulation at 200 Hz leads to a suppression of MEPs. <u>Bottom</u>: occurrence of modulation patterns with respect to mTh stimulation frequency. All patterns recorded in all muscles of 3 animals are included in the analysis (MK-RG 3 muscles, MK-AP 2 muscles, MK-MA 4 muscles). For all panels, data from all muscles and all animals shown in **Extended Data Figure 4**, stimulation parameters in **Supplementary Table 2**. For all panels, statistical significance was assessed with two-tail bootstrapping with Bonferroni correction: p<0.05 (*), p<0.01 (**), p<0.001(***).

First, we assessed the validity of H2 and H3. mTh stimulation alone at 50 Hz, delivered at the same amplitude used in the paired condition, did not elicit muscle responses (**Figure 3b**, **Extended Data Figure 4a)**, demonstrating absence of supra-threshold activation of direct projections to spinal motoneurons (either via the CST or other excitatory pathways). The absence of direct motor responses, however, does not rule out the possibility that spinal circuits are facilitated. We, therefore, tested whether mTh stimulation altered a purely spinal reflex response: the H-reflex. Peak-to-peak amplitudes of monosynaptic reflexes elicited by direct dorsal root stimulation (**Figure 1b**) were unchanged during continuous mTh stimulation at 50 Hz (**Figure 3c**, **Extended Data Figure 4b**), indicating that thalamic stimulation does not increase excitability of spinal circuits, either through suprathreshold or subthreshold modulation. Together, these two experiments expanded our previous findings^24^ and argued against H2 and H3.

Furthermore, to demonstrate that the effects were mediated by layer V corticospinal neurons, we surgically dissected the M1 hand region (**Figure 3d**, **Extended Data Figure 4c**): if the potentiation depends on these neurons, it should be abolished after dissection. As predicted, after M1 dissection, IC-evoked MEPs were no longer potentiated by concurrent mTh stimulation (**Figure 3d**, **Extended Data Figure 4d**), maximum percentage increase over muscles: MK-RG +161.4% and +10.9% and MK-AP +202.9% and +96.0% before and after disconnection, respectively), confirming H1. These combined controls demonstrate that facilitation occurs via a circuit loop through M1.

This implies that the modulation effects of corticospinal neurons must be mediated synaptically, and should, therefore, depend on the stimulation frequency, given the well-established frequency dependence of synaptic transmission and efficiency at sustained neural activations. For example, since excitatory postsynaptic potentials (EPSPs) are brief (5–20 ms), synaptically mediated excitatory inputs require a stimulation frequency high enough (>40 Hz) to sustain membrane depolarization^37,38^. Conversely, at high frequencies (>100 Hz), i.e., those clinically used in DBS, synaptic transmission becomes less efficient because of presynaptic mechanisms such as neurotransmitter depletion and presynaptic inhibition that limit the amplitude of EPSPs at high stimulation rates^39,40^. To test this, we paired IC stimulation at 1 Hz with mTh stimulation at different frequencies (30-200 Hz), while recording MEPs. We analyzed the MEP responses to individual IC stimulation pulses over time and observed distinct modulation patterns (**Figure 3e**). Specifically, we classified the modulation patterns of MEPs elicited by IC stimulation paired with mTh stimulation at different frequencies into three categories: (1) no potentiation (not observed), (2) sustained potentiation, that is the responses were consistently potentiated, (3) attenuation or suppression, as the response amplitudes progressively decreased after a few pulses. Note that the frequency of MEPs remained unchanged (1Hz), the only change is the frequency of mTh, therefore, this experiment tests the effects of mTh frequency on the potency of cortical modulation. As predicted, attenuation and suppression patterns were more prevalent with high frequency mTh stimulation (100-200 Hz, 40-100% occurrences over all animals and muscles), while low frequency mTh (30-80 Hz) reliably led to sustained potentiation (80-90% occurrences) (**Figure 3e**), further supporting H1 as well as providing guidance for tuning of stimulation parameters.

Overall, these results demonstrate that low-frequency mTh stimulation increases CST output both in animals with intact descending pathways and in animals with subacute and chronic CST lesions. Moreover, these electrophysiology experiments verified potential mechanisms of actions and allowed us to identify the optimal stimulation parameters: target and frequency. We next aimed at verifying whether this potentiation could lead to functional improvements of volitional motor control after CST lesions in awake behaving animals.

### Volitional arm and hand dexterity

Most daily activities require dexterous three-dimensional arm and finger movements, rather than the constrained single-joint movements assessed in simplified tasks. Thus, we designed a high-throughput experimental set-up to record arm and hand 3D kinematics while stimulating the mTh (**Figure 4a**). We then trained n=3 animals to perform a battery of upper-limb motor tasks, tailored to capture the stereotypical post-stroke motor deficits: 1) *touchscreen* task 2) *reach, grasp & pull* task, and 3) *constrained pinch* task.

**Figure 4:**
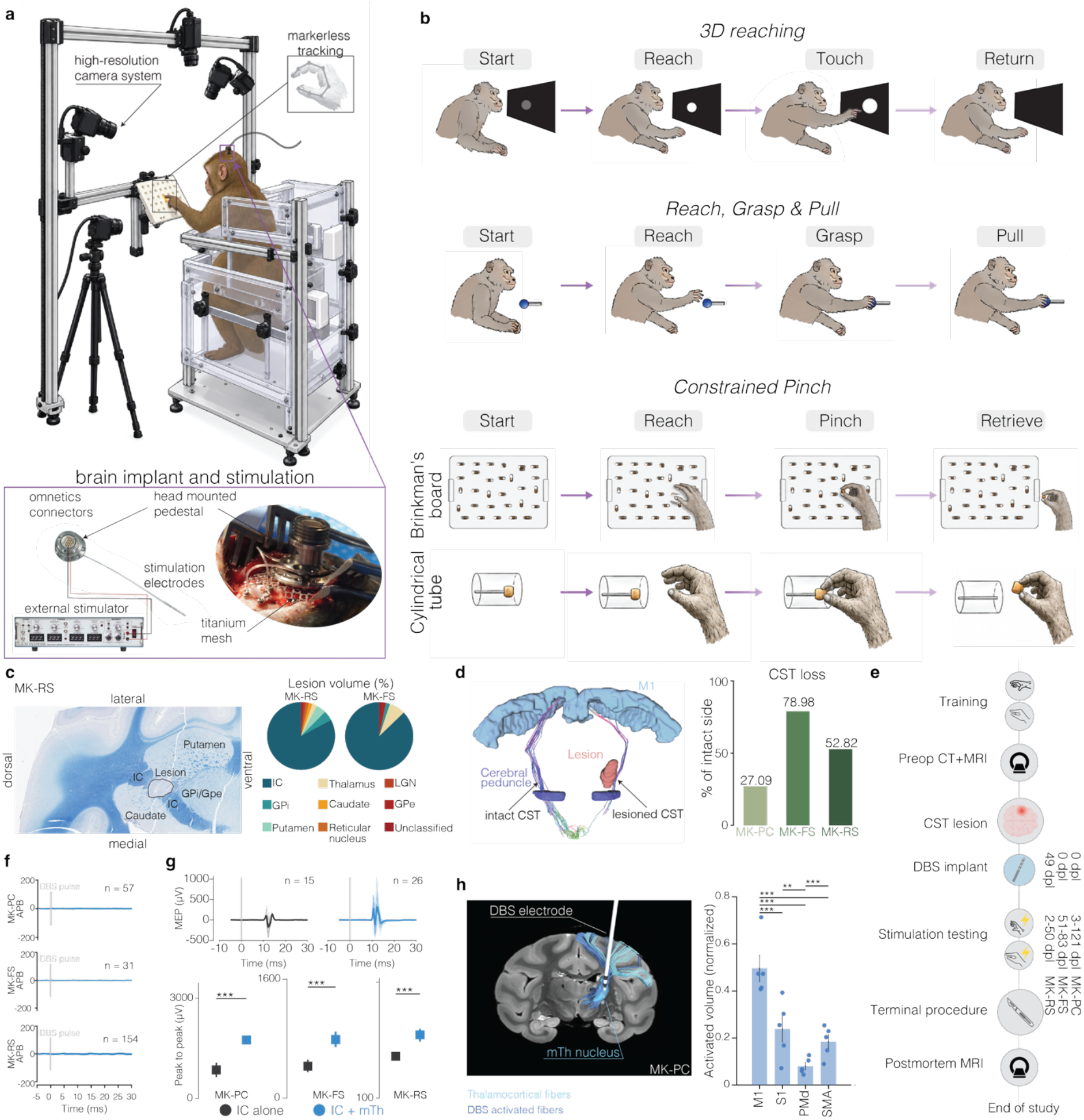
Experimental setup for chronically implanted animals and surgical procedures for chronic CST lesions and mTh DBS implant. (**a**) Experimental setup for recording 3D full-limb kinematics and behavioral performances in monkeys chronically implanted with a DBS lead. (**b**) Tasks designed to assess individual motor deficits. (**c**) example of HDFT of the CST in intact and lesioned hemisphere (MK-FS). On the right, bar plot of the volume loss of the CST in the lesioned side as a percentage of the CST volume in the intact side. (**d**) Histology analysis. <u>Left:</u> example of histology image, showing LFB-CV staining over a coronal brain section where the lesion is located. <u>Right:</u> pie charts showing the distribution of the lesion volume (in %) across multiple tissues for MK-RS and MK-FS. (**e**) Timeline for chronic experiments in n=3 monkeys, from task training to post-mortem imaging (‘dpl’=days post lesion). (**f**) examples of one muscle for all animals with mTh DBS alone at 50 Hz, used to confirm the location of the implant (**g**) <u>Top:</u> Example of MEP for MK-FS in a finger muscle (APB) with IC stimulation alone and IC stimulation paired with mTh DBS at low-frequency. <u>Bottom:</u> peak to peak MEP amplitude of one finger muscle for each animal to confirm implant location. Data from all animals and all muscles in **Extended Data Figure 5**. Data shown as mean ± 95% CI. (**h**) <u>Right:</u> example of 3D reconstruction of DBS implant within the mTh (blue nucleus). Thalamocortical (mTh to M1) and DBS activated (activation volume to M1) fibers reconstructed from HDFT are shown. <u>Left:</u> Bar plot showing the normalized volume of fibers recruited from the DBS and connected to different motor-related cortical regions (M1=primary motor cortex, S1=primary somatosensory cortex, PMd=dorsal premotor cortex, SMA=supplementary motor area). Each data point represents each implant for each animal (n=5). For all panels, statistical significance was assessed with two-tail bootstrapping with Bonferroni correction: p<0.05 (*), p<0.01 (**), p<0.001(***).

The *touchscreen* task was designed to evaluate the monkeys’ ability to perform an unconstrained three-dimensional reaching while supporting the weight of the arm (**Figure 4b**, *top row*). Animals had to reach from a starting position to a circular target displayed on a touchscreen monitor (positioned in front of the animal) and thereafter return to the starting position. After successful trials, ie., when the touch point was within the circular target, monkeys received a juice or food reward. In the *reach, grasp & pull task*, the animals were trained to reach for and grasp a spherical dynamometer and to pull it isometrically; a food reward was delivered when the exerted force crossed a predefined threshold that was adjusted daily and individually for each animal according to their recovery (**Figure 4b**, *second row*). Finally, in the *constrained pinch* task, animals were trained to retrieve and eat small food pellets from spatially constrained horizontal slots or from a narrow vertical cylindrical tube (**Figure 4b**, *bottom rows*), a previously described assay of finger dexterity^41,42^. Importantly, movement trajectories were not constrained, neither kinematically nor in time, in any of the tasks. Such design was intended to capture the coordinated arm extension, combined with hand grasping and individuated digit movements that characterize naturalistic every-day life upper-limb behaviors, without imposing a constrained timing or structure.

We then designed a set of behavioral and kinematic measurements to evaluate functional outcomes on task performances, strength and movement dexterity. Specifically, we quantified success rates, pulling forces, and markerless 3D kinematics of the full-limb and fingers^43–45^(see **Figure 4a** and Methods).

### Surgical protocol for CST lesion and DBS implant

Once animals reached a plateau performance in all the tasks, we induced a lesion of the CST by thermocoagulating the posterior limb of the internal capsule. Specifically, using the same robotic approach of the acute experiments we inserted a probe in the internal capsule, and we mapped its somatotopic organization by stimulating at different depths and recording MEP responses in different upper-limb muscles (**Extended Data Figure 5a**). We then thermocoagulated for 60 seconds at 85°C in the location with the lowest motor threshold and in a second location 2 mm apart. We confirmed the location and extent of the CST lesion by performing post-mortem histological analysis on dissected brains. We computed the percentage volume of the lesion invading the IC and surrounding tissues. In all monkeys, most of the lesion was successfully localized within the IC (87.2% and 82.8% for MK-FS and MK-RS, respectively) (**Figure 4c**), and post-mortem high-definition tractography confirmed that the lesion consistently reduced the volume of CST fibers compared to the intact hemisphere (percentage reduction: MK-PC 27.09%, MK-FS 78.98%, MK-RS 52.82%, **Figure 4d**).

During the same surgery, we chronically implanted a DBS probe in the mTh (**Figure 4e**). To confirm the location of the implantation, we 1) verified that mTh stimulation at 50 Hz did not directly activate CST fibers (**Figure 4f, Extended Data Figure 5b**), and 2) verified that low-frequency mTh stimulation led to a potentiation of arm and hand MEPs pre- and post-lesion (**Figure 1b**, **Figure 4g, Extended Data Figure 5c**). Finally, we confirmed the DBS lead location by co-registering post-operative CT with pre-operative MRI. Tractography combined with modeling of the stimulated tissue volume indicated that the fibers most recruited by the stimulation were those projecting to M1 (**Figure 4h** and **Extended Data Figure 5e**). Once the animals could move and climb freely in their housing and showed no signs of discomfort (2 to 5 days after injury, depending on the animal), animals were tested daily for up to 121 days post-lesion with one or two tasks per day to assess the evolution of the motor impairments after the lesion and the potential of the stimulation to immediately improve motor function, comparing stimulation ON and OFF in the same day (**Figure 4e**).

### Lesion of the CST results in chronic hemiparesis

After the lesion, the animals developed clear signs of hemiparesis, including loss of distal and proximal dexterity and strength.

Performances in the *touchscreen* task significantly dropped compared to baseline immediately after the lesion, with all monkeys achieving less than 5% of success rate the first week (5%, 0%, and 0% for MK-PC, MK-FS, and MK-RS, respectively) (**Figure 5a**). Performances then stabilized around 3-4 weeks post-lesion to values lower than pre-lesion, indicative of a permanent deficit (mean drop in success rate across week 3-10 compared to prelesion: -5% and -15% for MK-PC and MK-FS, respectively; MK-RS did not perform the task since the second week). A healthy unconstrained 3D reaching movement unfolds across two movement phases: reaching and stabilization phase. The reaching phase consists of the initial ballistic transport of the arm towards the monitor and relies primarily on coordinate proximal arm activation to generate a fast and smooth trajectory. As the hand approaches the target, control transitions to a stabilization phase, during which the animal must maintain a proximal and distal elevation against gravity while precisely regulating hand position to establish and maintain contact with the target (end-effector control). As expected, in intact animals, the reaching was fast and smooth, and the hand reached the target with minimal positional corrections and minimal loss of limb elevation (**Figure 5b**). To dissect which movement phase was mostly affected by the lesion, we computed a principal component analysis (PCA) of 38 kinematic parameters (**Supplementary Table 3**) capturing reaching speed, reaching smoothness, distal and proximal support, and end-effector control. In all animals, we identified two principal axes of separability between movements pre- and post-lesion (PC1-PC2 in MK-PC and MK-FS, and PC1-PC3 in MK-RS, **Figure 5c**) that explained most variance (72.6%, 73% and 70.1% of the data variability for MK-PC, MK-FS and MK-RS, respectively), and captured distinct dimensions of motor impairment: PC1 was primarily associated with the limb anti-gravity support and end-effector control features during the stabilization phase, whereas the second PC was predominantly associated with speed and smoothness during the ballistic reaching phase (MK-PC shown as example **Figure 5d, Extended Data Figure 6a** for the other animals). Individual kinematic features confirmed impairments in both the reaching and the stabilization phase. Indeed, we found that in all animals’ movements were slower (p<0.001) and less smooth (reduced number of velocity peaks, p<0.001, **Figure 5e**). Moreover, position variance, maximum positional displacement and vertical drop of the distal and proximal joints were significantly higher post-lesion (p<0.001, **Figure 5e**) confirming an impairment in the stabilization phase, which requires full-limb weight support and precise distal motor control and is strongly dependent on CST integrity. Importantly, movements in stroke survivors are usually slower, less smooth and more variable similarly to what is observed in our animals supporting the clinical validity of our injury model^5,46^.

**Figure 5:**
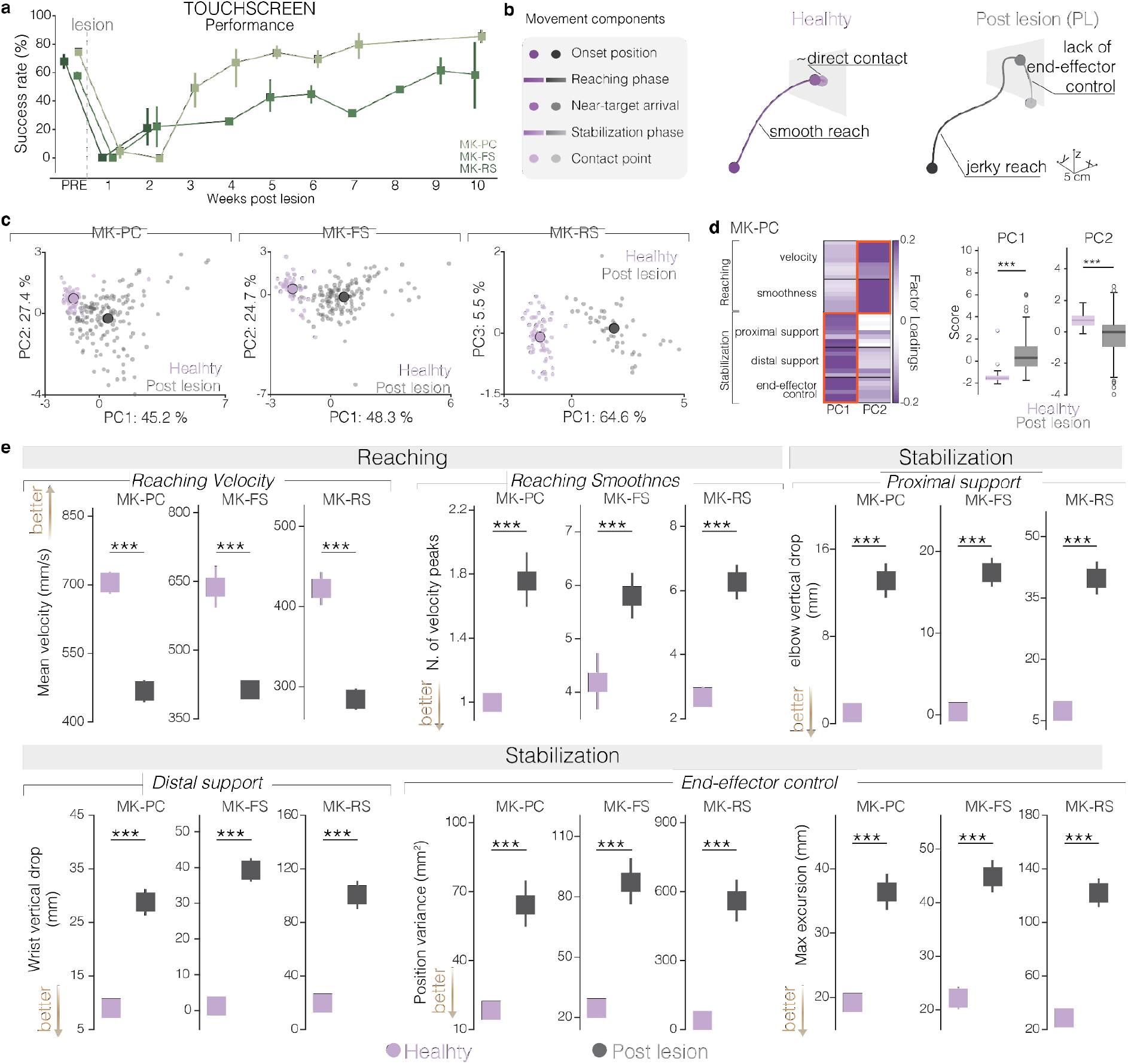
CST lesions impair reaching and stabilization in a 3D reaching task. (**a**) Task success rate over time. Data shown as mean across days ± 95% CI. (**b**) Example of wrist trajectories from MK-PC to highlight the movement components affected by the CST lesion. (**c**) PCA plane of the highest variance PCs that separate healthy and post-lesion movements. Each small dot is a reaching movement and the big dot is the mean across movements. (**d**) Loadings of features grouped according to their role in motor control. On the right, box-plot showing the PC scores before and after the lesion. For all boxplots, the whiskers extend to the maximum spread not considering outliers, central, top, and bottom lines represent median, 25th, and 75th percentile, respectively. Representative data for MK-PC, all data from all animals are in **Extended Data Figure 6a**. (**e**) Representative features for different motor control components, shown as mean across reaches ± 95%. For all panels, statistical significance was assessed with two-tail bootstrapping with Bonferroni correction: p<0.05 (*), p<0.01 (**), p<0.001(***).

We then evaluated strength and gross hand function with the *reach, grasp & pull* task. Immediately after the lesion, the animals were severely weaker. While strength (measured as the peak pulling force per trial) partially recovered over time (**Figure 6a**), animals still presented a significant residual deficit (drop in strength: -14%, -25%, -27% compared to healthy for MK-PC, MK-FS and MK-RS, respectively) in the chronic phase (i.e., > 4 weeks post-lesion, **Figure 6b**). Importantly, successful performance of this task requires more than sufficient force generation. Indeed, this task requires a goal-directed reaching component, during which the animal must appropriately pre-shape the hand, followed by the establishment of a stable grasp around the sphere before generating the pulling force (**Figure 6c**). Thus, successful execution depends on the coordinated control of proximal arm transport, distal hand configuration and subsequent force generation. Similarly to what we observed in the *touchscreen* task, the reaching phase of the movement was affected by the lesion both in speed and smoothness in all animals (**Extended Data Figure 6b**). Importantly, following the CST lesion, animals showed a significantly longer grasping time and higher end effector position variance, indicating that several attempts were required to successfully pull the sphere (**Figure 6d**). Within these attempts we observed two different failure modes: in some cases the animal was not able to successfully pre-shape the hand to grasp the sphere (that we refer to as pre-shaping fail); in other cases, instead, the animal could secure the grip but the force applied was not sufficient to cross the threshold (force fail) (**Figure 6e**). We quantified the relative contribution of these distinct failure modes by calculating the percentage of unsuccessful hand pre-shaping versus the percentage of unsuccessful forces per trial. Interestingly, after lesion the first behavior (pre-shaping failure) was prevalent (MK-PC: 62% vs 38%, MK-FS: 69% vs 31%, MK-RS: 59% vs 41%, percentage of pre-shaping failures versus force failures per trial), indicating that the deficits were not solely explained by residual weakness but also reflected a substantial impairment in hand motor control. This impairment is consistent with the critical contribution of the CST to the precise control of the hand and digits.

**Figure 6:**
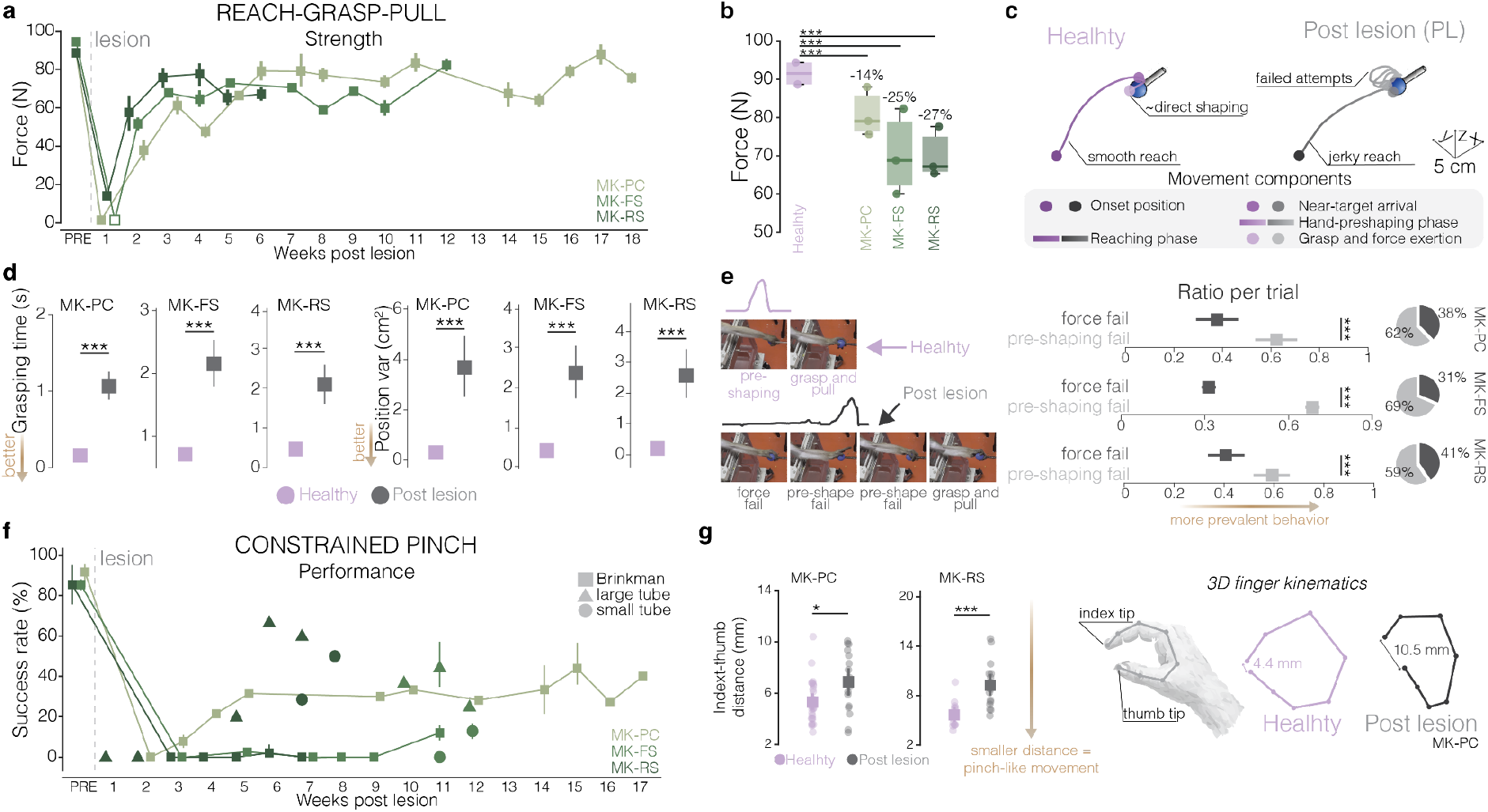
CST lesions impair strength and distal dexterity. (**a**) Maximum force exerted over time. Data shown as mean across days ± 95% CI. The white square indicates that the task was presented to the animal, but he was unable to perform it. (**b**) Residual strength deficit, calculated over the last 3 weeks of data collection of each animal. For all boxplots, the whiskers extend to the maximum spread not considering outliers, central, top, and bottom lines represent median, 25th, and 75th percentile, respectively. (**c**) Exemplary wrist trajectories from MK-PC to highlight the movement components affected by the CST lesion. (**d**) Grasping time and wrist position variance during the grasp. Data shown as mean across days ± 95% bootstrap CI. (**e**) <u>Left</u>: snapshots showing the development of pre-shaping and force deficits after the lesion (black), compared to healthy behavior (lilac). <u>Right</u>: Ratio per trial of the force failures and the pre-shape failures, for each animal. Data shown as mean across days ± 95% bootstrap CI and pie charts of the means. (**f**) Success rate over time for the *constrained pinch* task, with symbols highlighting the Brinkman Board or Cylindrical tubes. (**g**) Minimum distance between finger and thumb tip. On the right, an example of stick diagrams of the finger joints at the pinch frame for MK-PC, highlighting the increased distance after the lesion. For all panels, statistical significance was assessed with two-tail bootstrapping with Bonferroni correction: p<0.05 (*), p<0.01 (**), p<0.001(***).

Finally, we assessed individual digits control and hand dexterity using the *constrained pinch* task, either with the Brinkman board or the Cylindrical tubes. Successful retrieval from the horizontal and vertical holes required a high level of finger dexterity: animals had to coordinate multiple digits to grasp the food rather than relying on a gross, whole-hand grasp. Animals experienced a remarkable reduction in their ability to retrieve small food pellets, with success rate remaining substantially below healthy levels throughout the post-lesion period (**Figure 6f**). Specifically, MK-PC could not perform the task for the first two weeks post-lesion, and then retained a permanent moderate deficit never achieving above 50% success (up to 16 WPL), while MK-FS and MK-RS could never reliably perform the task (< 12%) and instead achieved moderate performances with the Cylindrical tubes (large cylinder: MK-FS 44.36%, MK-RS 66.66%; small cylinder: MK-FS 12.87%, MK-RS 50%). Importantly, although all animals retained the ability to produce movements of individual digits in some trials, they were unable to appropriately coordinate thumb and index movements to form a precise pinch. This deficit in inter-digit coordination is a hallmark of impaired hand dexterity and represents a major challenge for stroke survivors^5,47^. To quantify it, we reconstructed 3D finger kinematics of the thumb and index joints during food retrieval (**Figure 6g**). Following the lesion, we found a significant increase in the minimum thumb-index tip distance (MK-PC: 5.31 vs 6.89 mm p<0.05, MK-RS: 5.68 vs 9.27 mm p<0.001, mean distance in healthy vs post-lesion, **Figure 6g**), indicative of an abnormal thumb-index configuration and impaired formation of the functional pinch posture. These findings indicate that CST injury disrupted the precise inter-digit coordination required to organize the fingers into a functional pinch posture.

In summary, animals presented deficits in *i*) the ballistic phase of the reaching, *ii*) the end-effector stabilization, *iii*) strength, *iv*) whole-hand prehension, and *v*) digits coordination.

### Systematic calibration of DBS parameters

After implantation, we conducted a systematic procedure to fine-tune the stimulation parameters to immediately facilitate (and not induce) voluntary movements across the different motor tasks.

For each animal and implant, the stimulation was first turned ON at 50 Hz using the electrode configuration that yielded maximum MEPs potentiation during the implantation surgery. We then increased the amplitude until the animals showed signs of perceived discomfort or possible paresthesia, identified as abrupt movements on the chair; at that point, we lowered the amplitude by 0.2 mA. Stimulation amplitude was further adjusted by observing immediate changes in motor performance during the different tasks with the stimulation ON. The final amplitude satisfied two constraints: (i) it was lower than the threshold for perceived discomfort, and (ii) it was lower than the amplitude identified during the implantation surgery as eliciting direct CST activation. This procedure was repeated daily, and the amplitude typically varied by less than 1 mA across sessions (**Extended Data Figure 7a**). Pulse width was also adjusted based on animal discomfort and immediate behavioral effects: for MK-FS and MK-PC the pulse width was always 200uS whereas for MK-RS we used 200uS 91% of the time and 400uS 9% of the time. Crucially, all amplitudes and pulse widths were within the range of clinically approved DBS parameters and stimulation was always delivered at 50 Hz. Finally, we periodically tested other electrode configurations to check for potential changes in the efficacy of the stimulation. For each new configuration, amplitude was ramped up starting from 0 mA as described above. In MK-PC the electrode configuration remained unchanged in 100% of the recordings, in MK-FS in 94% of the recordings, and in MK-RS in 82% of the recordings (**Extended Data Figure 7a**).

### DBS restores movements otherwise impossible

We next assessed the effects of mTh DBS at low-frequency on the different tasks. Animals were tested daily with one or two tasks per day repeated with stimulation ON and OFF each day. Outcomes were computed for each animal independently and compared between DBS ON and DBS OFF. We first asked whether the stimulation could improve task performance during the period of most severe impairments, i.e., in the first 1-2 weeks after the lesion when animals were not able to reach or grasp. Because animals could not perform the dexterity tasks until several weeks after the lesion, we focused this first analysis on the *touchscreen* and *reach, grasp & pull* tasks. Moreover, analysis was performed only for MK-PC and MK-RS because in MK-FS the electrode moved during the first days, when impairments were most severe; stimulation was restored from day 51 post-lesion after replacement of the lead (see Methods).

During the period of greatest impairment, without stimulation, animals were unable to reach towards the touchscreen, despite repeatedly attempting the movement (**Figure 7a**). Interestingly, when turning the DBS ON, both animals were able to generate sufficient elbow extension and arm support to contact the monitor as early as 2 days post-lesion (DPL) (**Figure 7a**). Specifically, MK-RS successfully completed (i.e., managed to perform a full reach and hence contact the monitor) 83% of the attempted reaches with DBS ON, and 0% with DBS OFF. Similarly, MK-PC successfully completed 85% of the reaches with DBS ON, and only 55% with DBS OFF (**Figure 7a**). A few days later, once the reaching component could be performed, both animals with low-frequency mTh-DBS achieved a higher task success rate in the *touchscreen* task (**Figure 7b**) (MK-PC performed 20% vs 62.5% and MK-RS performed 12.08% vs 28.93%, reported as mean across days (7 and 8 DPL and 7 and 15 DPL) with DBS OFF and ON, respectively). Similar severe impairments were observed in the first days in the *reach, grasp & pull* task (**Figure 7c**). We analyzed the first day in which animals attempted to perform the task (3 and 5 DPL for MK-PC and MK-RS, respectively). Without stimulation, animals consistently failed to complete the grasp required to initiate the pull (**Figure 7c-d**), resulting in a 0% task success rate. In contrast, with DBS ON, animals were able to successfully grasp and pull, reaching a 50% task success rate (**Figure 7e**). Moreover, even among the few trials in which animals were able to produce force, without stimulation, the peak pulling force applied remained low, never exceeding 15 N (average of the peak forces across trials with DBS OFF: 5.96N for MK-PC and 14.15N for MK-RS). Stimulation immediately increased force production resulting in a substantially higher peak force (average across trials with DBS ON: 50.77N for MK-PC and 28.22N for MK-RS) (**Figure 7f**).

**Figure 7:**
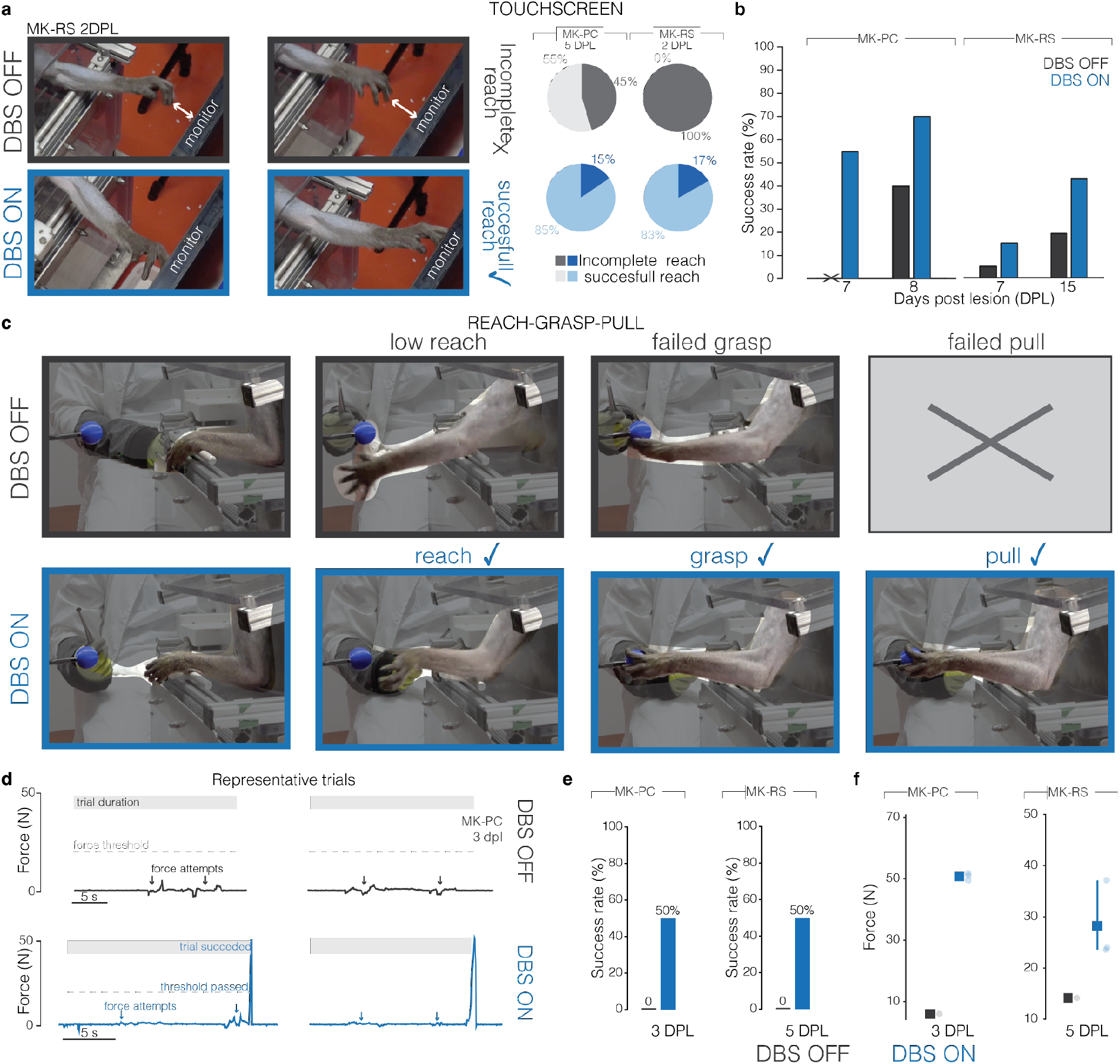
DBS improves motor function immediately after the lesion, allowing for movement otherwise impossible. (**a**) <u>Left</u>: Snapshot from the 2nd day post lesion of MK-RS showing attempted reaches, that can successfully be completed on the touchscreen monitor only with DBS ON. <u>Center</u>: pie charts of incomplete and successful (i.e., the animal contacts the touchscreen) reaches for MK-PC and MK-RS on the first day of tuned DBS. (**b**) Success rate without and with DBS for the first dates where successful trials were performed. (**c**) Snapshots from MK-PC performing the pull task immediately after the lesion. Without DBS, monkeys could not grasp and pull the sphere, or they could only occasionally exert low forces. With DBS ON the animals could reach, grasp, and pull. (**d**) Example of representative force trace showing successful pull exertion only when DBS was turned ON. (**e**) Bar plot of success rate in the pull task, for the first day on which the animals performed the task without and with DBS. (**f**) Maximum force exerted during pull trials, shown as mean across trials ± 95% CI.

Overall, these results show that DBS facilitated gross arm and hand movements otherwise impossible. We next sought to explore whether DBS could have a beneficial effect also in the chronic phase, after partial spontaneous recovery, when residual deficits persisted.

### Motor thalamus DBS improves distal anti-gravity support and end-effector stabilization

We next assessed effects of mTh stimulation at low-frequency on all the trained behavioral tasks throughout the entire testing period, which extended up to 4 months after the lesion. Daily, we compared DBS ON and OFF. The order of the two conditions was pseudorandomized across sessions to control for possible effects of order or fatigue within a session. The comparison between DBS ON and OFF was always performed using the optimized stimulation parameters of each animal; sessions in which amplitude, frequency or electrode configuration were varied were used as control conditions and analyzed separately.

In the *touchscreen* task, all animals showed an immediate increased task success rate when the stimulation was turned ON across multiple sessions (**Figure 8a**). We then explored which components of motor control accounted for this improvement. mTh-DBS produced a moderate enhancement of the reaching phase. Specifically, reaching movements were significantly faster in one animal (MK-FS) and smoother in two animals (MK-PC and MK-RS) when the stimulation was switched ON (**Extended Data Figure 7b**). Instead, in all animals, DBS reliably improved multiple kinematics features reflecting distal (i.e., wrist) anti-gravity support and end-effector control during the stabilization phase (**Figure 8b, Extended Data Figure 7b**). Specifically, MK-PC and MK-FS significantly improved in 86% of the calculated features related to distal weight-support and 83% of the features related to end-effector control, whereas MK-RS improved in 100% of features within these two domains (**Figure 8c**). The robustness of the benefit provided by DBS during the stabilization phase of an unconstrained 3D reaching, is further highlighted by the fact that, when performing a PCA with DBS OFF and DBS ON trials, for all animals the first PC, which accounts for most variance of the data and significantly separates the two conditions, was purely dominated by stabilization features (**Extended Data Figure 7c**). In other words, stabilization-related features were the primary contributors to the separation between DBS ON and OFF trials, highlighting end-effector control as a key component of the kinematic improvements associated with DBS. Functionally, restoration of distal support and end-effector control resulted in reduced hand drops and more precise target contacts (**Figure 8d**).

**Figure 8:**
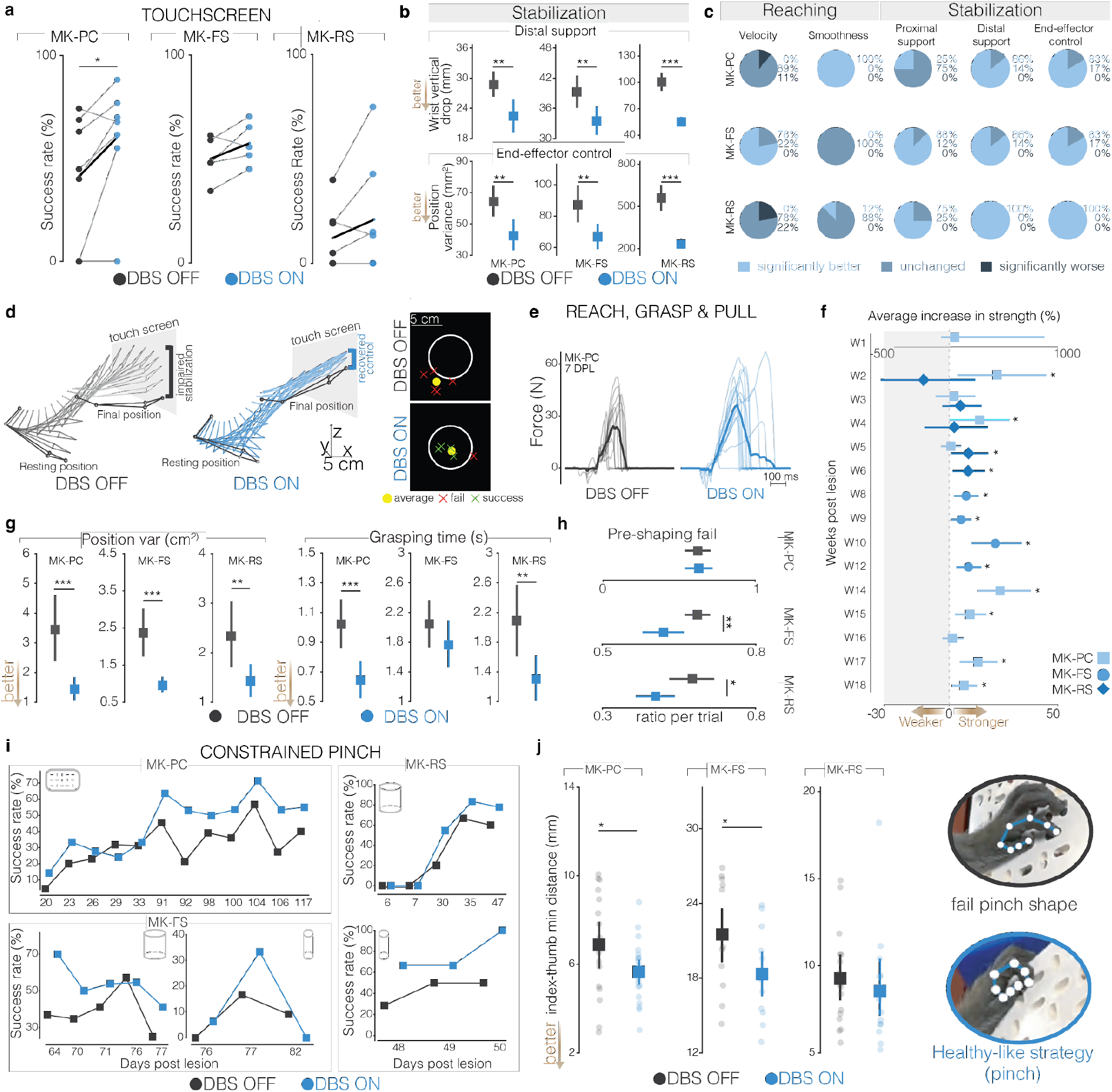
DBS stimulation improves end-effector control, strength and dexterity. (**a**) Success rate of *touchscreen* task for all days, bold line represents the mean across days. (**b**) Representative stabilization kinematic features, shown as mean across trials for multiple days ± 95% CI. (**c**) Pie charts showing the percentage of features belonging to each movement component that improved significantly, did not change or worsened significantly with DBS. While reaching features present more inconsistent improvements, the benefits of DBS in the stabilization phase were robust over animals. (**d**) Example of stick diagram of the arm joints during a reaching, highlighting the stabilization impairment without DBS. On the left, example of a *touchscreen* target (white circle) and contact locations separated for success and failed trials. (**e**) Example of force traces without and with DBS. The bold line represents the average across trials. (**f**) Percentage increase in strength with DBS ON (compared to DBS OFF) for all animals across weeks. Data shows as mean across sessions ± 95%CI. (**g**) Position variance of the wrist and grasping time during the *reach, grasp & pull* task, without and with DBS. (**h**) Ratio per trial of the pre-shape failures with DBS ON and OFF, for each animal. Data shown as mean across days ± 95%CI and pie charts of the means. (**i**) Success rate over time for the *constrained pinch* task, separated according to the method used (Brinkman Board or cylindrical tubes). (**j**) Minimum distance between finger and thumb tip, each point represents a pinch movement. On the right, a snapshot highlighting an incorrect pinch frame (characterized by high fingertips distance) and a corrected pinch achieved thanks to DBS. For all panels, statistical significance was assessed with two-tail bootstrapping with Bonferroni correction: p<0.05 (*), p<0.01 (**), p<0.001(***).

### DBS effects were specific to mTh and low-frequency stimulation

In dedicated recording sessions we compared the influence on task performance of three stimulation parameters: i) stimulation frequency, to test whether the electrophysiology evidence from anesthetized monkeys would also expand to motor behavior; ii) stimulation amplitude, to confirm that the effects observed were amplitude-dependent and that an optimized configuration existed; and iii) stimulation location, to confirm the spatial specificity of mTh-DBS. In the *touchscreen* task, the success rate was higher when the DBS was delivered at 50-80 Hz (20 and 30% improvement from baseline) compared to 130 Hz, i.e., the frequency conventionally used in clinical DBS (10%, data from MK-RS 13 DPL) (**Extended Data Figure 8a**). This result was consistent with the suppression of MEPs observed at high-frequencies in our electrophysiology experiments (**Figure 3e**). Additionally, task performance dropped when the stimulation amplitude was not properly tuned (either too low to produce a detectable effect or too high, potentially causing discomfort) or when the stimulation was delivered at contacts located outside of mTh (**Extended Data Figure 8a**). Finally, we verified that the stimulation was not inducing undesired movements, as demonstrated by no significant differences in the resting position with and without DBS (**Extended Data Figure 8b**). This suggests that indeed the stimulation was inducing immediate facilitation of movements and not imposing an overt motor response, and that the facilitation was specific to low-frequency stimulation within the mTh.

### Motor thalamus DBS improves strength and hand gross motor function

We next assessed whether mTh DBS increased strength, measured as the peak pulling force per trial. Across several weeks of testing, animals generated greater pulling forces with stimulation ON up to +55% (**Figure 8e-f**), resulting in improved task success rate (**Extended Data Figure 7d**). Moreover, the stimulation reduced position variance and grasping time, indicating that fewer attempts were required to secure the grasp (**Figure 8g**). We then evaluated whether the stimulation improved the ability to secure hand grip. Immediately, when the stimulation was turned ON, the percentage of pre-shaping failures per trial was significantly reduced, indicating improved hand positioning and grasp formation (**Figure 8h**). Finally, analysis of the reaching portion of the *reach, grasp & pull* task reproduced the dichotomy observed in the *touchscreen* task: with stimulation ON, reaching movements were smoother in only two animals, whereas end-effector control improved in all three (**Extended Data Figure 7e**).

Together, these results confirmed that mTh DBS improves distal and end-effector control, as well as gross hand motor function, by facilitating successful grasp formation and increasing pulling force. Similarly to our previous findings in the *touchscreen* task, these effects were specific to mTh and low-frequency stimulation (**Extended Data Figure 8c**).

### Motor thalamus DBS improves hand dexterity

Finally, we asked whether DBS could also improve the more dexterous aspects of hand function, which remain a major challenge for individuals recovering from stroke for which current neuromodulation approaches have shown limited efficacy. Across multiple testing days, in all animals, DBS ON consistently resulted in higher pellet-retrieval rates compared to DBS OFF (in MK-PC performance was better with stimulation 10/11 days, in MK-FS 4/6 days and in MK-RS 6/6 days, **Figure 8i**).

Importantly, kinematic analysis revealed a significant reduction in the minimum thumb–index finger distance when the stimulation was ON (MK-PC 6.89 vs 5.64 mm, p<0.05, MK-FS 21.51 vs 18.32 mm p<0.05, MK-RS 9.27 vs 8.55 mm p>0.05, average min distance with DBS OFF and ON, respectively, across trials **Figure 8j**), suggesting that the improvement in retrieval success was associated with improved finger dexterity during the pinch. Similarly to the other tasks, these effects were specific to mTh stimulation at low-frequency (**Extended Data Figure 8d**).

Together, these findings indicate that, beyond improving proximal and gross hand motor function, mTh stimulation can also facilitate the precise inter-digit coordination required for dexterous object retrieval.

## Discussion

Here we showed that continuous stimulation of the motor thalamus improved strength, distal and end-effector control and, most importantly, fine motor control of the hand in three monkeys with chronic CST lesions. These effects were specific to low-frequency stimulation and were visible immediately when turning ON the stimulation. Moreover, the observed movements were not induced by an overt motor response generated by the stimulation: the animals retained volitional control, which was amplified by DBS. We here discuss the implications of our results for the understanding of motor recovery and for future clinical applications.

### CST lesions reproduce post-stroke weakness and loss of dexterity

Humans with stroke develop hallmark motor symptoms of hemiparesis, including persistent weakness and loss of proximal and distal dexterity. Mounting evidence in patients suggests that these motor deficits emerge primarily from the loss of corticospinal axons rather than from cortical damage per se^48^. However, in humans this has largely been inferred from correlations between lesion anatomy, corticospinal excitability and impairment, and cannot be tested causally. We, therefore, first developed and characterized a non-human primate model of subcortical stroke, via a controlled, electrophysiology-guided lesion of the posterior limb of the internal capsule. We reliably reproduced the hemiparetic phenotype in three monkeys. Specifically, lesion of the CST resulted in deficits across multiple domains of motor control, including *i*) reaching velocity and smoothness; *ii*) pulling strength; *iii*) end-effector control and hand gross motor function; and *iv*) finger individuation, that closely resembled human post-stroke motor impairments. Interestingly, the most severe deficits in grasping time and inter-digit coordination appeared to be associated with greater CST loss and larger lesion volume, whereas reaching speed, smoothness, and proximal support did not show the same pattern. Nevertheless, these deficits persisted over time in all animals, allowing us to test our approach in both the acute and the chronic phase (over 4 months post-lesion). Consistent with the corticospinal origin of these deficits, existing primate models based on ibotenic acid injections over the motor cortex, which damage the cortex while largely sparing the descending axons, produce more limited dexterity deficits that often recover within weeks^49^. Therefore, in addition to demonstrating causally that CST disconnection is sufficient to produce post-stroke weakness and loss of dexterity, our model represents a technical advance over previous approaches.

### Facilitation of CST output requires connections to corticospinal neurons are necessary to improve dexterity and hand functions

We then hypothesized that motor deficits resulting from CST damage could be immediately alleviated by augmenting corticospinal drive to spinal motoneurons. While direct stimulation of corticospinal axons would result in an overt motor response that necessitates a precise delivery of the stimulation, a facilitation of CST output would instead integrate with ongoing cortical dynamics, preserving volitional control even with continuous stimulation^50^. The monosynaptic glutamatergic connections from the motor thalamus to layer V pyramidal neurons of the motor cortex (i.e., corticospinal neurons) provide an ideal native synaptic interface for this purpose. We found that low-frequency continuous stimulation of the motor thalamus produced immediate improvements in distal motor control, such as end-effector stabilization, grasp formation, and inter-digit coordination, alongside improvements in strength. Notably, the stimulation produced no involuntary movements. Importantly, the effects were specific to low-frequency stimulation, and CST output potentiation was abolished after M1 dissection supporting a synaptically mediated mechanism.

The electrophysiological and behavioral outcomes of mTh-DBS suggest that the synaptic position of stimulation relative to corticospinal neurons is an important determinant of its assistive efficacy. Indeed, CST output potentiation was greater when stimulation was targeted within the motor thalamus rather than its afferents (i.e., the dentate nucleus or globus pallidus internus). Previous research aligns with our findings: in 12 chronic post-stroke individuals, dentate nucleus DBS, whose excitatory projections reach the motor cortex disynaptically via the ventrolateral thalamus, produced therapeutic changes but no assistive motor effects and only limited increase in MEP amplitude^18^. The only other reports of immediate MEP potentiation, similar to our results, came from M1 stimulation conditioned by premotor cortex stimulation^51,52^, and the corresponding behavioural effect was a reduction in grasp duration^53^. Yet from these results it was not clear whether the stimulation improved configuration of the digits during manipulation like in our thalamic approach. This could possibly be explained by the fact that corticocortical synapses produce EPSPs roughly ten times smaller than those of thalamocortical synapses^54^, suggesting that immediate improvements in hand dexterity require a large increase in corticospinal drive to spinal motoneurons.

Finally, the effects of mTh stimulation were largest for distal support, end-effector control and inter-digit coordination compared to reaching speed, smoothness, and proximal support. If assistive effects require a monosynaptic connection to corticospinal neurons, as our comparison across stimulation targets suggests, these functions would be improved precisely because they are driven by CST. Reaching and proximal functions, instead, may be controlled, and may recover, through corticoreticular, corticorubral and corticopontine projections, which mTh stimulation cannot facilitate as effectively because of the disynaptic connection between the motor thalamus and the reticulospinal neurons originating in the brainstem. This interpretation remains a hypothesis that would need to be tested with selective manipulation of the individual pathways, but it offers a possible framework for the open debate on the role of different descending pathways in motor recovery after stroke^22,55^.

### A clinically accessible approach to immediate motor restoration

The translational potential of this approach rests on the combination of a clinically validated procedure and an immediate functional effect. DBS is an established technology, and the motor thalamus is a standard target for essential tremor. Additionally, our results were obtained with clinically approved DBS leads and parameters, possibly promoting a rapid translation of our approach. Indeed, the stimulation configurations used in this study could be implemented directly with commercially available DBS systems, and stimulation parameters remained stable over time. Only stimulation amplitude was slightly adjusted daily, comparable to the adjustments patients routinely perform themselves with commercial DBS systems for other applications^56^.

A further advantage is that stimulation produced immediate improvements in motor performance without requiring the detection of movement intention, or continuous adaptation of stimulation parameters to neural signals, or pairing with training sessions. This assistive effect could provide a direct behavioral readout for selecting stimulation settings, potentially allowing parameters to be optimized according to the motor functions most impaired in each patient. Such immediate assistance may also complement rehabilitation by enabling patients to perform movements that would otherwise be difficult^57^ although whether it enhances long-term recovery remains unknown. Determining the relationship between immediate motor facilitation and lasting therapeutic benefit will be an important next step toward clinical translation.

Finally, while previous studies using functional electrical stimulation of the muscles (FES) or spinal cord stimulation (SCS) reported improvements in hand motor functions, such as grasp and object transfer^58–60^, these approaches often require different stimulation locations and configurations for different movements. In our approach, instead, the stimulation engages the native thalamocortical interface of corticospinal neurons, which lie within the same cortical territory regardless of whether they control proximal or distal muscles. A single configuration was therefore sufficient to enable 3D reaching movements as well as gross and fine hand motor functions, and the same configuration potentiated MEPs in arm, hand and finger muscles. Because one configuration improved both proximal and distal function, parameter selection in patients would not require separate settings for different movements, which could simplify clinical use.

### Limitations

A few limitations of our study need to be acknowledged. First, the sample size is small (n=3 for the behavioural experiments, n=7 for the electrophysiological experiments), but it is consistent with the principle of the 3Rs and comparable to previous non-human primate studies^28,61–63^. The thermoablation of the internal capsule does not reproduce the full pathophysiology of stroke, including the inflammatory processes that follow an ischemic or hemorrhagic event. Yet, thermoablation allows reproducible lesions limiting the number of animals necessary as imposed by the principle of the 3Rs. Moreover, the similarity of the deficits reported in our study and in a previous single case^64^ with those observed clinically support the validity of the model. Yet we did not test effects of the stimulation on spasticity due to the absence of a reliable test in non-human patients. Finally, because all animals received stimulation throughout the testing period, we cannot exclude that repeated use of DBS influenced the spontaneous evolution of the deficits over the four months of testing, and we therefore focused our analyses on the immediate, assistive effects of the stimulation.

### Future directions

Future studies in human patients will now need to answer these questions in a larger population. Moreover, in humans the motor thalamus comprises three nuclei with distinct preferential projection targets: the ventral anterior (VA), ventral oral posterior (VOP), and ventral intermediate (VIM) nuclei^30^. Thalamocortical fibers originating in the VIM/VOP connect to M1, and those in the VA to the premotor cortex. We could hypothesize that if our theory of the need of a monosynaptic connection to corticospinal neurons is correct, the VIM/VOP would represent the optimal nucleus within the motor thalamus. Future studies will need to confirm this and whether mTh stimulation could help for other functions requiring fine motor control such as speech and swallowing. Our recent preliminary results point to this direction^65^, although it remains unclear whether upper-limb and orofacial functions could be addressed with a single DBS system. Finally, we recently showed that post-stroke residual CST sculpted the strength of spinal reflexes produced by each pulse of SCS to promote functionally relevant muscle activation patterns^50^. We, therefore, hypothesize that a combinatorial approach in which DBS is used to increase CST output and SCS is then applied to further empower those descending signals might promote a synergistic effect of the two systems separately and so stronger behavioral benefits.

## Methods

### Animals involved in the research

All animal procedures were approved by the University of Pittsburgh Institutional Animal Care and Use Committee (protocol number IS00025637). We utilized a total of 7 *Macaca Fascicularis* (5 males: MK-PC 4.8Kg, MK-FS 6.2Kg, MK-RS 5.4Kg, MK-AP 5.0Kg, MK-RG 4.8Kg, 1 female: MK-KA 6.5Kg) and 1 *Macaca Mulatta* (1 male: MK-MA, 6.0 kg); all animals ranged from 4 to 8 y.o. The animals were housed in the Division of Laboratory Animal Resources at the University of Pittsburgh. When possible, the animals were pair-housed with another animal prior to any surgical procedure. The animals were not water- or food-restricted and were given daily enrichments (novel food, toys, puzzles). Three animals (MK-PC, MK-FS, MK-RS) received a chronic DBS implant and a lesion of the CST to induce chronic hemiparesis, after being trained on upper-limb and hand tasks. All animals (except MK-RS) underwent terminal procedures during which we performed electrophysiology experiments (**Figure 1b**). Unfortunately, MK-RS was euthanized before the study could be completed due to a severe infection of a spinal array implanted at day 50 post-lesion for another project, in respect of the 3R rules and in agreement with the endpoints in our veterinary authorization. Detailed information on which animals were involved in specific experimental procedures is reported in **Supplementary Table 1**.

### Preoperative imaging

For all animals, we acquired pre-surgical images to plan the DBS implant and terminal procedures. During the first imaging session, we acquired T1-weighted magnetic resonance imaging (MRI) scans of the brain (400 μm isotropic, TR/TE: 6000/3.7 ms, 7T Siemens whole body human system). MRI was performed under full anesthesia induced with ketamine (10 mg/kg, i.m.) and maintained under isoflurane (1-3%, 2L/min, inhalant). During the second imaging session, we first performed a survival surgical procedure in which we implanted five fiducial screws (1.5 mm × 3.5 mm, KLS Martin) in the skull at non-coplanar depths. We then acquired a computed tomography (CT) scan of the skull with the implanted screws (250 μm isotropic, Epica Vimago GT30). The fiducial screws allowed registration to the surgical robot (see below). This approach has been described in detail previously^31^. The procedure was performed using standard aseptic techniques, under full anesthesia induced with ketamine (10 mg/kg, i.m.) and maintained under isoflurane (1–3%, 2 L/min, inhalant). Animals then received carprofen (Rimadyl, 4 mg/kg, i.m., once daily for 3 days) and enrofloxacin (Baytril, 2.5 mg/kg, i.m., once daily for 5 days). Both imaging procedures were performed with the animals secured in a customized plastic stereotactic frame to facilitate co-registration of the images^66^.

### Surgical protocol for terminal surgeries

All animals, except MK-RS, underwent terminal procedures with electrophysiological experiments. These procedures were performed under full anesthesia, induced with ketamine (10 mg/kg, i.m.) and maintained under continuous intravenous infusion of propofol (1.8–5.4 ml/kg/h) and fentanyl (0.2–1.7 ml/kg/h), which were preferred over isoflurane as they minimally affect electrophysiological responses (Toossi, A. et al. *J. Neural Eng.,* 2019). Heart rate, oxygen saturation, end-tidal CO2 and body temperature were monitored continuously throughout the procedures. During each terminal surgery, we performed the following procedures: 1) intramuscular needle electrode implantation, 2) robot-assisted deep probe implantation, 3) subdural ECoG implantation 4) epidural dorsal root implantation. Certified neurosurgeons performed these surgical procedures (Drs. Jorge Gonzalez-Martinez, Peter Gerszten, UPMC, Pittsburgh, USA). At the end of the experiments, the animals were euthanized with a single injection of pentobarbital (86 mg/kg) and perfused with 4% paraformaldehyde (1 L/kg) for further tissue imaging.

#### Implantation of intramuscular needle electrodes

Electromyographic (EMG) activity was measured with 13mm percutaneous electrode needles placed on the main upper limb and hand muscles, specifically Left Bicep (BI), Tricep (TRI), Extensor carpi radialis (ECR), Extensor Digitorum Communis (EDC), Flexor Carpi Radialis (FCR), Flexor Digitorum Communis (FDC), Abductor Pollicis Brevis (APB), and Flexor Digiti Minimi (FDM). All EMG data was amplified, digitally processed, and recorded using the Ripple Neuro Grapevine and Trellis software at a sampling frequency of 30,000 Hz.

#### Implantation of deep probes

We used the ROSA One(R) Robot Assistance Platform (Rosa robot) to perform implantation of deep probes, following a workflow similar to the one used clinically for stereo-EEG (SEEG) and DBS implantation in patients. Prior to the surgery, we co-registered MRI and CT images using the ROSA One Brain Application. We then planned implantation trajectories for the desired targets. Coordinates are reported relative with the vertical axis defined by the AC-PC plane, and the anterior-posterior position scaled to the AC-PC distance of each animal to account for differences in brain size. Specifically, to access the corticospinal tract (CST) we defined a first trajectory targeting the hand area of the CST within the posterior limb of the IC in the right hemisphere (stereotactic coordinates anterior-posterior: in the middle of AC-PC, vertical: 0 mm, lateral: +8 mm). To deliver stimulation to the ventral lateral thalamic nucleus (that we here refer to as the motor thalamus, mTh; corresponding to the VIM/VOP target used clinically) we planned a second trajectory (stereotactic coordinates anterior-posterior: length AC-PC distance/4 +0.5 mm, vertical: 0 mm, lateral: +4 mm). Additionally, in MK-AP and MK-RG we also inserted a stimulation probe in the ipsilateral Globus Pallidus Internus (GPi, stereotactic coordinates anterior-posterior: length AC-PC distance/2, vertical: -2.5 mm, lateral: +11 mm) and in the contralateral Dentate Nucleus (DN, stereotactic coordinates anterior-posterior: length -8.5mm, vertical: -8 mm, lateral: -3.5 mm), whereas MK-MA received only the DN implant. The targets for all trajectories were defined on the MRI imaging and validated by a trained functional neurosurgeon, avoiding vasculature and the ventricles. During the procedure, each monkey was positioned prone in a stereotaxic head frame (Kopf, Model 1530, Tujunga, CA, USA), and the previously implanted fiducial screws were used to register the ROSA robot position with respect to the animal. The robot was then used to implant the probes. During terminal procedures, all electrodes targeting either mTh, DN and GPi were Dixi 8-electrode leads (DIXI Microdeep® SEEG Electrodes), secured on the skull at the insertion point with a bolt. IC stimulation was delivered either through a DIXI 8-electrode lead or via 2 radiofrequency cannulas (Abbott medical) implanted 5mm apart in the posterior limb of the internal capsule. DN, GPi and mTh stimulation was tuned by ramping up amplitudes at 50Hz and observing for direct muscle activation or undesired responses such as increased heart rate. We ultimately chose the amplitude value at which a plateau in the IC induced MEP peak-to-peak was observed. Continuous stimulation of the 3 nuclei was then paired with IC stimulation at 1Hz to verify the presence of MEPs potentiation, and the amplitudes were further adjusted in the tolerated range defined above to find the value inducing highest potentiation. During the experiments, IC stimulation was delivered as either single pulses or bursts of cathodal, charge-balanced, symmetric square pulses. In all cases, we first identified the threshold at which we could measure consistent muscle activity from at least 2 muscles, the stimulation probes were moved along the trajectories to minimize such threshold. Stimulation was delivered via an AM stimulator (model 2100 A-M Systems, Sequim, WA, USA), with the parameters listed in **Supplementary Table 2**. Final position of DN, GPi, IC and mTh probes during terminal experiments was confirmed by post-mortem imaging and by inducing small thermoablation around the electrodes used for stimulation (Neuro N-50 Lesion Generator, Inomed Medizintechnik, Germany)

#### M1 stimulation and recording

After implantation of the deep probes, we performed a 20 mm diameter craniotomy over the central sulcus of the left hemisphere and removed the dura to expose the primary motor cortex (M1). Functional motor areas of the arm were identified through anatomical landmarks and intra-surgical micro-stimulation. We positioned a 6-electrode ECoG array (6 contact platinum subdural electrode, AD-TECH Medical Instrument Corporation, Oak Creek, WI) so that one contact covered the identified position of the hand knob of M1. We then tuned ECoG stimulation to elicit muscle activity on hand and arm muscles (MK-AP: 3mA burst at 400Hz, 25ms duration at 0.5Hz, 500µs pulse width, and MK-RG: 7mA burst at 300Hz, 25ms duration at 0.5Hz, 300µs pulse width). Stimulation was delivered in a bipolar configuration, using the contact over M1 as cathode and the most proximal contact as anode. In order to record from the same array, we connected the ECoG to the Ripple Grapevine system (sampling frequency of 30,000 Hz). During the recordings, a third channel, laying posteriorly to M1 but still on cortical tissue was used as ground and reference.

#### Epidural dorsal root stimulation experiments

Epidural dorsal root stimulation experiments were conducted in MK-RS and MK-KA. In MK-RS, spinal cord was exposed and a laminectomy was performed at C5-C7 vertebrae. An 8-contact spinal cord lead (Medtronic Vectris) was inserted between T1 and T2 and slid in position covering the left dorsal aspects of the spinal cord between level C5 and T1. In MK-KA, we performed a larger laminectomy at C3-T1 level and we used a bipolar probe to deliver stimulation at level C6-C7 where motoneurons controlling hand muscles are located. We used either 1Hz (MK-RS, 1Hz, 200uS pulse width, 2.2mA) or 50 Hz burst epidural dorsal root stimulation (MK-KA, 1s burst at 50 Hz, 0.9mA 200uS pulse width) to induce muscle activation. We then paired epidural dorsal root stimulation with mTh stimulation at 50Hz at the amplitude used during IC related experiments. We then compare either peak-to-peak or area under the curve (AUC) in the two conditions (**Figure 3c, Extended Data Figure 3c**).

### Data analysis of electrophysiology experiments

#### Analysis of cortical evoked potentials

To analyze cortical evoked potentials elicited by 1 Hz stimulation of the mTh, DN or GPi, we recorded local field potentials in the M1 subdural ECoG strip. Data was filtered offline with a band-pass Butterworth 2nd order filter (20-1000 Hz) and a Notch filter (59-61 Hz) to remove noise interference. From each stimulation pulse, we extracted 75ms epochs (5ms before and 70ms after the stimulus). Each epoch was baseline-corrected by subtracting the mean of the pre-stimulus interval, and we then calculated the peak-to-peak amplitude of the evoked potential in each epoch. Stimulation-triggered averages were computed for visualization.

#### Analysis of muscle activity

Electromyographic activity was bandpass-filtered (10–500 Hz) and notch-filtered (59–61 Hz) using second-order Butterworth filters to remove artifacts and line noise. To measure peak-to-peak amplitudes of motor evoked potentials (MEPs), we extracted 35-ms epochs around each stimulus (5ms before to 30ms after). When the stimulation eliciting the muscle response was a burst (M1 stimulation, or dorsal root stimulation in Mk-KA) and no clear MEP was elicited, we instead calculated the area under the curve of the rectified EMG as an amplitude measure, similar to previous studies that quantified EMG responses to stimulation trains (Overduin et al., 2012 *Neuron*; Barra et al., 2022; Ho et al., 2024).

#### Analysis of frequency-dependent modulation of muscle responses

To assess the effect of mTh stimulation frequency, we performed experiments in which we paired IC stimulation at 1-2 Hz with mTh stimulation at 30, 50, 80, 100, 130 or 200 Hz. We visually inspected the peak-to-peak amplitude of MEPs over time in all conditions and classified them into four categories: “no potentiation”, “sustained potentiation”, “attenuation”, “suppression”. “Attenuation” and “suppression” applied to patterns in which the peak-to-peak amplitude of subsequent responses was progressively reduced and abruptly abolished, respectively. “Sustained potentiation” referred to patterns in which pairing IC stimulation with mTh stimulation amplified the MEP without any temporal modulation. “No potentiation” applied to patterns in which no observable change in the evoked responses occurred. To validate the classification, we computed the ratio between the peak-to-peak amplitude of the last and first 20 pulses. Thus, a ratio lower than 1 indicates a progressive decrease of the MEP responses, suggesting attenuation and/or suppression.

### Chronic DBS implant and lesion survival surgical procedure

Three animals (MK-PC, MK-FS and MK-RS) underwent chronic CST lesion and mTh DBS implant, and performed behavioral experiments up to 4 months after lesion. The lesion of the CST and chronic implant of the DBS electrodes in the motor thalamus were performed in the same procedure by a trained functional neurosurgeon in standard sterile conditions. These procedures were performed under full anesthesia induced with ketamine (10 mg/kg, i.m.) and maintained under continuous intravenous infusion of propofol (1.8–5.4 ml/kg/h) and fentanyl (0.2–1.7 ml/kg/h). Similar to the terminal procedures, we used the ROSA One(R) Robot Assistance Platform (Rosa robot) for both implant of chronic DBS probes and to perform accurate lesion of hand and arm fibers in the posterior limb of the internal capsule (Ho et al, iEEE Proceedings, 2022). We first co-registered MRI and CT images using the ROSA One Brain Application; we then planned 2 trajectories: one targeting the hand area of the corticospinal tract (CST) within the IC at the AC-PC level, and another targeting the mTh, both in the right brain hemisphere. We planned the trajectories of the probe in the ROSA software using the same coordinates reported above, avoiding vasculature and the ventricle, and all the plannings were validated by a neurosurgeon. During the procedure, each monkey was positioned prone in a stereotaxic head frame (Kopf, Model 1530, Tujunga, CA, USA), and the implanted fiducial screws were used to register the ROSA robot to the monkey. Electromyographic activity was measured with 13mm percutaneous electrode needles placed on the main upper limb and hand muscles, specifically Left Bicep, Tricep, Extensor Carpi Radiali, Flexor Carpi Radiali, Abductor Pollicis Brevis, and Flexor Digiti Minimi. All EMG data was amplified, digitally processed, and recorded using the Ripple Neuro Grapevine and Trellis software at a sampling frequency of 30,000 Hz.

In all the 3 animals, we first implanted the chronic DBS lead targeting the mTh. For that, we designed an ad hoc implant with either a 4-channel Dixi lead (DIXI Microdeep® SEEG Electrodes) or a 4-channel Medtronic DBS lead (Medtronic, Minneapolis, USA). The leads were cut to the desired length (10-15 cm), and every single wire was soldered to a 16-channel Micro 360 Circular Omnetics connector (Omnetics Connector Corporation, Minneapolis, USA). The connectors were then encapsulated in silicone (734 flowable sealant) and fixed within a titanium pedestal (Blackrock Microsystems, USA). In order to stabilize the implant on the skull, a titanium mesh (TiMesh, Medtronic) was modelled on the 3D-printed skull for each monkey, and a footplate was attached to it using 5 screws (Buri SA, La Chaux-de-Fonds, Switzerland). Mesh, footplate, and screws were ETO sterilized individually and assembled in a sterile environment before being sterilized again pre-operatively to limit implant-related risk of infection. During the procedure, we first fixed the mesh and footplate with self-drilling fiducial screws (1.5 mm × 3.5 mm, KLS Martin) and then proceeded with the robot-guided drilling of penetration holes at the entry point for the mTh trajectory. In MK-FS and MK-RS, the implant of the DBS electrode along the desired trajectory was driven by a slotted cannula (AD-TECH, USA), whereas for MK-PC, we relied on a slotted bolt at the entry point for the insertion of the mTh lead. In both implant strategies, depth was defined based on pre-surgical planning. After insertion, the pedestal was loosely fixated on the footplate, and the IC probe (TCBA011, RF TC Brain Electrode, Inomed Medizintechnik, Germany) was temporarily implanted following a similar robot-guided procedure. Hand-specific CST projections were identified by mapping electromyographic response along the insertion trajectory within the posterior limb of the IC. Specifically, stimulation was delivered at 1Hz between 0.2 and 2 mA, and the location with the lowest threshold of activation was deemed the hotspot for hand muscle activity recruitment (±2 mm from planned target in all animals). We then used the IC stimulation to verify the location of the mTh electrodes. For this, we paired 1Hz suprathreshold stimulation of the IC with mTh continuous stimulation at 50-80Hz. We then compared different configurations of mTh stimulation for all the animals to find the one that maximized muscle recruitment compared to IC stimulation alone. We also further verified that mTh stimulation alone was not inducing any muscle activation by ramping amplitudes from 0.2mA to 5.0mA at 50Hz. We then performed 2 thermocoagulations of the CST fibers within the IC at the hotspot location and 2 mm apart, each for 1 minute at 85℃ with a radiofrequency generator (Neuro N-50 Lesion Generator, Inomed Medizintechnik, Germany). Lastly, the mTh electrode’s pedestal was fixed on the footplate permanently, and dogbones were used to secure in position the electrode’s wire from the insertion point to the pedestal. The IC probe was removed, and the skin incision was sutured. Correct positioning of the mTh implant was verified with post-operative CT scan after co-registration with pre-operative MRI imaging.

### Replacement surgeries

Over the course of the study, the implant was replaced twice in MK-PC (at 16- and 84-days post-lesion, DPL) and once in MK-FS (at 49 DPL). As part of our routine post-implantation verification, a CT scan acquired after the first implant in MK-PC showed that the lead was positioned slightly posterior to the planned trajectory. To optimize targeting of thalamocortical fibers, we elected to reposition the lead, which increased the estimated volume of cortical projections recruited by the optimal electrode configuration by ∼10%. The position of the second implant was confirmed with a CT scan at 27 DPL (**Figure 4h** and **Extended Data Figure 5d**). At 43 DPL, we observed muscle twitches when ramping up stimulation amplitude that had not been present during implantation or during previous recording sessions. We hypothesized that the sEEG lead had gradually shifted closer to the internal capsule, favoring direct recruitment of corticospinal fibers, and the implant was therefore replaced at 84 DPL. In MK-FS, a CT scan at 22 DPL indicated that the lead was not at the intended target; behavioral data were collected without stimulation during this period, and the implant was replaced at 49 DPL. All subsequent implants were performed using DBS leads (Medtronic), used in clinical practice and they did not show any signs of overtime lead migration. All replacement surgeries followed the procedure described above. The titanium mesh and footplate were left in place and reused to secure the pedestal of any new implant.

### Design and behavioural training of upper-limb tasks

MK-PC, MK-FS, and MK-RS were equipped with a metallic collar and trained to sit on a plexiglass and metal chair (GrayMatter, US) designed to restrain movement of their right arm and ensure a stable body position during left arm movements. All the 3 animals were trained to execute the *touchscreen* task, the *reach, grasp & pull* task, the *Brinkman Board* task, and a constrained pinch task as explained in section “*Volitional arm and hand dexterity”*. Food pellets, fruits, and fruit juice were used as positive rewards, and additional food was provided after training to complete daily dietary requirements. In all the tasks, the relative position between the animal and the task items (touchscreen, dynamometer, Brinkman’s board and pinch cylinders) was set before baseline collection based on animal size and maintained constant throughout all data acquisition. In all the tasks, animals were trained to start and stop each reach from a resting bar steadily fixated on the primate chair.

#### Touchscreen task

The touchscreen task was implemented using an ad-hoc Python script based on PsychoPy libraries^67^. In this task, the animal had to reach from a starting position (i.e., located mid-way between the center of the touch screen (ELO ET1902L LCD, Elo Touch Solutions, USA) and the acromion, at the same height) each of 9 targets (arranged in a clock-like fashion, placed 15 cm from its center) on a touchscreen (positioned in front of the animal) and move back from each target to the starting position. During each repetition of task, an acoustic trigger accompanied the appearance of a white circular target (radius 33-45 mm, adapted to each animal’s skills on that task) on the screen and, in case of success, acoustic feedback on a different tone was played. Targets were chosen randomly among 9 different positions (positions) for at least 3 repetitions per position during healthy recordings. No acoustic feedback was played for failed reaches.

#### Reach, grasp & pull

During the *reach, grasp & pull* task, visual feedback on the exerted force and success threshold was visible to the animal; success and trial start were tied to the same acoustic feedback as the touchscreen task. We recorded pull force with a force gauge (Mark 10, model m5-20, capacity 100N) and 3D-printed personalized attachments in the form of a small sphere and a large sphere (radius ∼2.40 cm and ∼3.23cm respectively). The force threshold was adjusted for each animal everyday according to their recovery, in order to keep the animals engaged and motivated.

#### Brinkman Board

In the Brinkman’s board task the animals were asked to retrieve food pellets from small openings (6mm deep, 14mm long and 7mm wide) on a 3D printed board (22cm length, 12cm wide and 1.2cm thick x 120), as in^42,68^. The board was angled at 45 degrees and placed centered in front of their impaired arm.

Pellets were approximately circular in 2mm±1mm of radius. Animals were given a minute of time per condition to retrieve and eat as many food pellets as possible.

### Constrained Pinch

Animals were trained to retrieve a piece of fruit from within a vertical plexiglass cylinder, presented in front of the tested hand at shoulder level. Two cylinder sizes were used, 3 cm (small) and 6.5 cm (large) in diameter, which restricted the space available for hand opening and forced the animal to shape its hand and use a pinch grip (Sasaki et al., 2004; Alstermark et al., 2011). Each session consisted of a maximum 3 minutes of consecutive pinches.

### Behavioural data acquisition and experimental recordings

Once the animals were trained, we collected baseline recordings of all tasks before the lesion for at least 15 days and more than 3 recording sessions per task. Following the surgical CST lesion, animals started data recording between 2-5 days post-implantation, depending on their recovery. MK-FS, which implant has to be substituted, was recorded without stimulation until day 49 post lesion. MK-PC performed 42 sessions over a 4-month period, MK-FS 35 sessions over a 3-month period, and MK-RS 30 sessions over a 2-month period. All sessions lasted less than 3 hours. During each session, the animal was required to perform one or two of the trained tasks without stimulation and with DBS ON. During some sessions, the same task was repeated with different parameters to either optimize the stimulation or as a control. Stimulation of the mTh was provided using an AM stimulator (model 2100 A-M Systems, Sequim, WA, USA). Details on parameters used and electrode configurations across sessions are reported in **Extended Data Figure 7a.**

During behavioural experiments, animals were filmed with 4 high-resolution Sony Handycam (model FDR-AX43) cameras. Cameras were manually started, and video recordings were synchronized using LEDs signals triggered from a DAQ (USB6009, National instruments, USA) and securely placed in the experimental area. A trigger was sent at the beginning and end of each recording and at the start of each trial during the *reach, grasp & pull* and *touchscreen* tasks. Videos were recorded at 120 frames/second.

We used the Ripple Neuro Grapevine and Trellis software to record pull forces during the reach, grasp & pull task at a sampling frequency of 30,000 Hz. We also recorded analog signals with the trigger, the pull force, and the stimulation pulses from the AM stimulator. We also used an ad-hoc Python script to record touchscreen performances during the *touchscreen* tasks. Particularly, from every repetition we logged target position, touch position, time of touch (w.r.t the appearance of the target).

### Reconstruction of 3D upper limb kinematics

We used DeepLabCut (DLC) and Anipose to extract 3D pose estimation of the animals’ upper limb and hand joints^43–45^. Before each recording, we recorded calibration videos where a Charuco board was shown on the camera field of views, moving in space for around 90 seconds to ensure optimal calibration. We then used DLC (network ResNet -101) to label anatomical landmarks and reconstruct 2D pose estimation of at least two cameras for each task. Each DLC model was trained on a healthy dataset of manually labelled frames, and further optimized by adding monkey-specific labelled samples. We trained 2 models for each animal, the first for arm tracking during the *touchscreen task* and the *reach, grasp and pull* task, and the latter on finger and hand anatomical landmarks to study hand shape and finger movements for the *Brinkman’s board task* and *constrained pinch task*. The first model tracked shoulder, elbow, wrist (ulnar and radial) and index finger, whereas the second model was trained to identify index and thumb fingertip, joint and knuckle, and webbing point (see **Figure 6g**). We then used Anipose to reconstruct 3D joint position by combining multiple cameras tracking and the calibration files. Further details on the optimization and tracking procedure are published in a previous work from our lab (Uribe et al. 2025 *iEEE*).

### Analysis of task performances

For each recording day, task performance was quantified separately for each behavioral task. Daily performances were calculated separately for the DBS OFF and DBS ON conditions, with DBS ON trials recorded using optimal stimulation parameters (or control parameters, **Extended Data Figure 8**). When a condition was tested in multiple consecutive blocks, these blocks were averaged to calculate a single daily success rate. For the DBS OFF condition, only blocks recorded consecutively within the same session were included to minimize potential confounding effects of fatigue or carryover effects from previous stimulation. In the *touchscreen task*, performance was assessed as the success rate, defined as the proportion of trials in which the monkey touched the target within its boundaries. In the *reach, grasp & pull task*, performance was assessed by computing the strength (calculated as the maximum force value in Newton exerted by the monkey during the trial) and the success rate, defined as the proportion of trials in which the monkey exerted a sufficient force to pass the predefined threshold. Finally, in the *constrained pinch* task, for the success rate we counted the number of food pellets that were extracted and brought to the mouth and divided it by the total number of food pellets that the monkey attempted to retrieve.

### Analysis of kinematic performances

For each task, we computed 3D kinematic analysis to study the influence of the CST lesion and the stimulation of motor control on a subset of sessions, ensuring that at least 2 days of recording were included. For each task, we used a custom-designed user interface to manually label events of interest in the video recordings. In the *touchscreen task*, we defined the movement performed as composed of two different phases: reaching and stabilization. The identification of the reaching phase was done by marking the moment in which the left hand left the metallic bar to when the maximum extension was reached, hence the finger was near the monitor. The stabilization phase was considered to start right after the reach phase and conclude once the touch contact was finalized (as identified by the monitor changing to the success or the failure state). For each of this phase, we computed kinematic variables, reported and detailed in **Supplementary Table 3**. On this large set of features, we then computed PCA. Before performing PCA analysis, features were centered to have mean 0 and scaled to have standard deviation of 1 (MATLAB2024a). We grouped the features according to the motor control component they represent, specifically: reaching velocity (that is maximum and mean velocity of all joints during the reaching phase), reaching smoothness (that is number of velocity peaks and log dimensionless jerk for all joints during the reaching phase), proximal support (that is relative and absolute vertical drop and drift of the shoulder and elbow during the stabilization phase), distal support (that is relative and absolute vertical drop and drift of the wrist during the stabilization phase), end-effector control (that is variance, excursion and speed during the stabilization for the wrist marker). In the *reach, grasp & pull task*, we defined the movement performed as composed of two different phases: reaching and pre-shaping/grasping. The first phase was identified as starting when the hand of the monkey left the resting position to the last frame of the video before contact with the dynamometer sphere was made. The second phase started hereafter and concluded when the hand of the monkey was secured around the sphere, muscle contraction was visible and force was exerted. During the reaching phase, we calculated the smoothness (as the log dimensionless jerk of the wrist) and during the pre-shaping and grasping phase we computed the position variance and the grasping time (as in **Supplementary Data Table 3**). Finally, for the constrained pinch, we marked two events: first, the starting of the movement as the moment in which the animal started to interact with the food (marked by the finger touching the edge of the Brinkman slots or the cylindrical tubes) and secondly, the end of the movement as the moment in which the food was securely pinched and removed from its location. We computed the minimum Euclidean distance between the thumb and index tip during this window as a measure of finger dexterity in pinch posture.

### Post-mortem imaging analysis

#### Reconstruction of deep electrodes

To confirm the location of deep electrodes (mTh, DN, GPi) implanted during terminal procedures to perform electrophysiology experiments, we acquired post-mortem T1-weighted MPRAGE MRI sequences. The implant tract was identified by visual inspection. Images were then co-registered across animals using anatomical landmarks and overlaid. The mean stimulation-site location across animals was calculated in FIJI, with coordinates referenced to the center of each implant tract.

#### High-definition fiber tracking

Following perfusion, the brain was dissected for post-mortem imaging and immersed in 1× PBS for 48 hours, followed by at least 1 week in PBS+ 0.2% gadolinium (1 mmol/mL, Gadavist, Bayer). The specimen was then placed in a custom 3D-printed holder filled with Fomblin to match tissue magnetic susceptibility and prevent dehydration during scanning. MRI data were acquired using a 9.4 T, 31-cm horizontal-bore Bruker AV3 HD animal scanner equipped with a high-performance 12-cm gradient set (maximum gradient strength: 660 mT/m) and a 72-mm quadrature birdcage RF coil. The imaging protocol included T2-weighted MRI (125 μm isotropic resolution; TR/TE = 1500/60.57ms) and diffusion-weighted MRI (0.25 mm isotropic resolution; b-values of 1800, 3400, and 5200 s/mm², with 30, 40, and 90 diffusion directions, respectively, and six b = 0 images). Diffusion tensor estimation and tractography were performed using DSI Studio (http://dsi-studio.labsolver.org). The accuracy of the diffusion gradient table (b-table) orientation was assessed by comparing reconstructed fiber orientations with those of a population-averaged template. Diffusion data were reconstructed using generalized q-sampling imaging with a diffusion sampling length ratio of 0.4 (0.6 for MK-RS). A deterministic fiber tracking algorithm^69^ was used with augmented tracking strategies^70^ to improve reproducibility. To quantify the damaged corticospinal tract, we used the following parameters: anisotropy threshold was randomly selected between 0.5 and 0.7 otsu threshold, angular threshold was randomly selected from 45 degrees to 90 degrees, step size was set to voxel spacing, tracks with length shorter than 30.0 or longer than 300.0 mm were discarded, and the tract-to-voxel ratio was set to 1.0. To reconstruct the CST, a first ROI was placed over the primary motor cortex, and another ROI was manually drawn at the cerebral peduncle. The tracts were manually pruned to exclusively include CST fibers. Tractography statistics (specifically total volume in mm^3) were saved directly from DSI Studio.

#### Histology

To confirm the location and extent of the corticospinal tract (CST) lesion, we performed post-mortem histological analysis of the brains of MK-RS and MK-FS. Post-mortem MRI was first used to localize the internal capsule (IC) lesion, which guided the manual dissection of a 5mm rostrocaudal tissue block centered on the lesion. Each block was cut into 10µm coronal sections, yielding approximately 500 sections per animal. Luxol Fast Blue with Cresyl Violet (LFB-CV) was used to stain each slice and visualize myelinated white matter and neuronal cell bodies. For each stain, we analyzed a series of 25 sections spaced 200µm apart, using adjacent, non-overlapping series. Stained sections were imaged at 20× magnification, and the lesion and the main anatomical structures (see **Figure 4c**) were manually delineated using image analysis software (QuPath). Finally, a custom Python script was used to compute the volume of the lesion overlapping each labeled structure, as reported in **Figure 4c**.

### Statistical analysis

All data analyses and figure generation were conducted in MATLAB R2024a (MathWorks, Natick, MA, USA). In all box plots, the central line indicates the median, while the lower and upper boundaries of the box correspond to the 25th and 75th percentiles, respectively. Whiskers indicate the data range excluding outliers. Scatter plots display the mean together with 95% bootstrap confidence intervals, estimated using 10,000 resampling iterations. If more than 15 samples were present, outliers were removed using MATLAB’s *rmoutliers* function. For statistical comparisons involving more than three observations, we used a non-parametric bootstrap approach to estimate uncertainty without assuming a specific underlying data distribution. For each comparison, 10000 bootstrap replicates were generated by randomly sampling the observed data with replacement. The difference in means between the groups was calculated for each replicate, yielding an empirical distribution of mean differences. Statistical significance was assessed using a two-sided test with a significance threshold of (alpha=0.05), based on the corresponding 95% confidence interval. When multiple pairwise comparisons were conducted, the significance threshold was adjusted using the Bonferroni correction. P-values were estimated by approximating the bootstrap distribution of mean differences with a Gaussian distribution and calculating the two-tailed probability associated with a difference at least as extreme as zero.

## Authors contribution

E.P., A.D., N.M., and M.C. conceived the study. E.P. and J.G.M secured funding. A.D., N.M., E.M.G., M.C., J.G.M., and E.P. designed the experiments. L.L., L.T., C.J. and K.J. performed the image and histology analysis. Z.X. developed the software to integrate the deep lab cut data for the kinematic analysis with the help of M.U., and processed the data with the help of A.D., I.N.. A.D., N.M., J.G.M., and E.P. designed the neurosurgical approach and J.G.M. and P.G. performed the surgery. K.H. provided anesthesia support during all procedures. L.T. and Z.X. assisted with all monkey surgeries. A.D. and N.M. collected the data during the anesthetized experiments, with the help of C.G., L.L., E.P., and M.C.. A.D. and N.M. developed the experimental rig and designed the tasks for behavioral experiments.. A.D., N.M. and T.S. built the electrodes. A.D., N.M., E.M.G., C.J., and K.H. trained the animals and collected behavioral data. A.D. and N.M. analyzed the data and created the figures. E.P., J.G.M. and M.C. helped with data interpretation. E.P., A.D. and N.M. wrote the manuscript and all authors contributed to its editing.

## Declaration of interest

The authors declare no competing interests.

### Acknowledgments

We would like to thank Evan Rogers for his help during the recording sessions of MK-PC, Kevin Hitchen for providing support for postmortem imaging, Kelly Puglisi and the Pitt Biospecimen Core for providing support for the histology analysis. We also thank the University of Pittsburgh DLAR for their support with animal care. We wish to thank Zimmer Biomet for lending the ROSA robot to use in our surgeries (we declare no conflict of interest). LL was supported by a scholarship from the Dompé Foundation. NM was supported by the Swiss national science foundation postdoc mobility grant. The study was executed through the support of internal funding from the Department of Physical Medicine and Rehabilitation at the University of Pittsburgh to E.P. and from the Department of Neurological Surgery at the University of Pittsburgh to M.C. and J.G.M. Additionally, E.P. and M.C. disclose support for the research described in this study from the Walter L. Copeland Foundation and J.G.M. disclose support for the research described in this study from the Hamot Health foundation. Finally, this work was supported by the National Institutes of Health (R01NS131428 to E.P. and J.G.M. and R01NS122927 to J.G.M.).

## EXTENDED DATA FIGURES

**Extended Data Figure 1:**
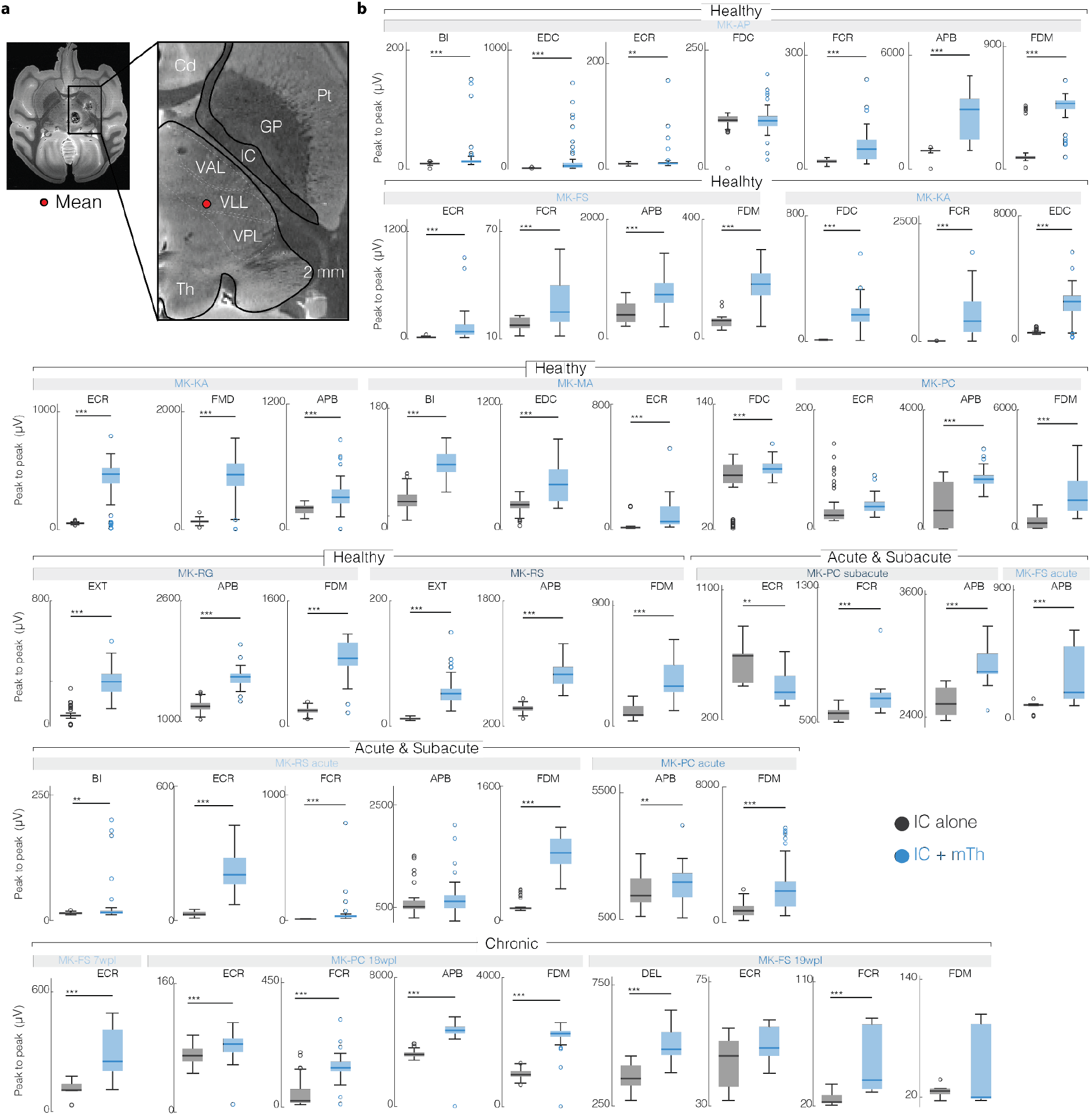
Potentiation of MEPs with mTh stimulation in healthy animals and animals with acute and chronic CST lesions. (**a**) MRI showing reconstructed mTh stimulation location, shown as average across animals. White dotted line shows the anatomical border of the motor thalamus and its nuclei. (**b**) For all muscles and animals, boxplots of the peak-to-peak amplitudes of MEPs with IC stimulation alone (black) and IC stimulation paired with low-frequency stimulation of the mTh (blue). We report only muscles that showed responses to stimulation. Stimulation parameters are reported in Supplementary Table 2.

**Extended Data Figure 2:**
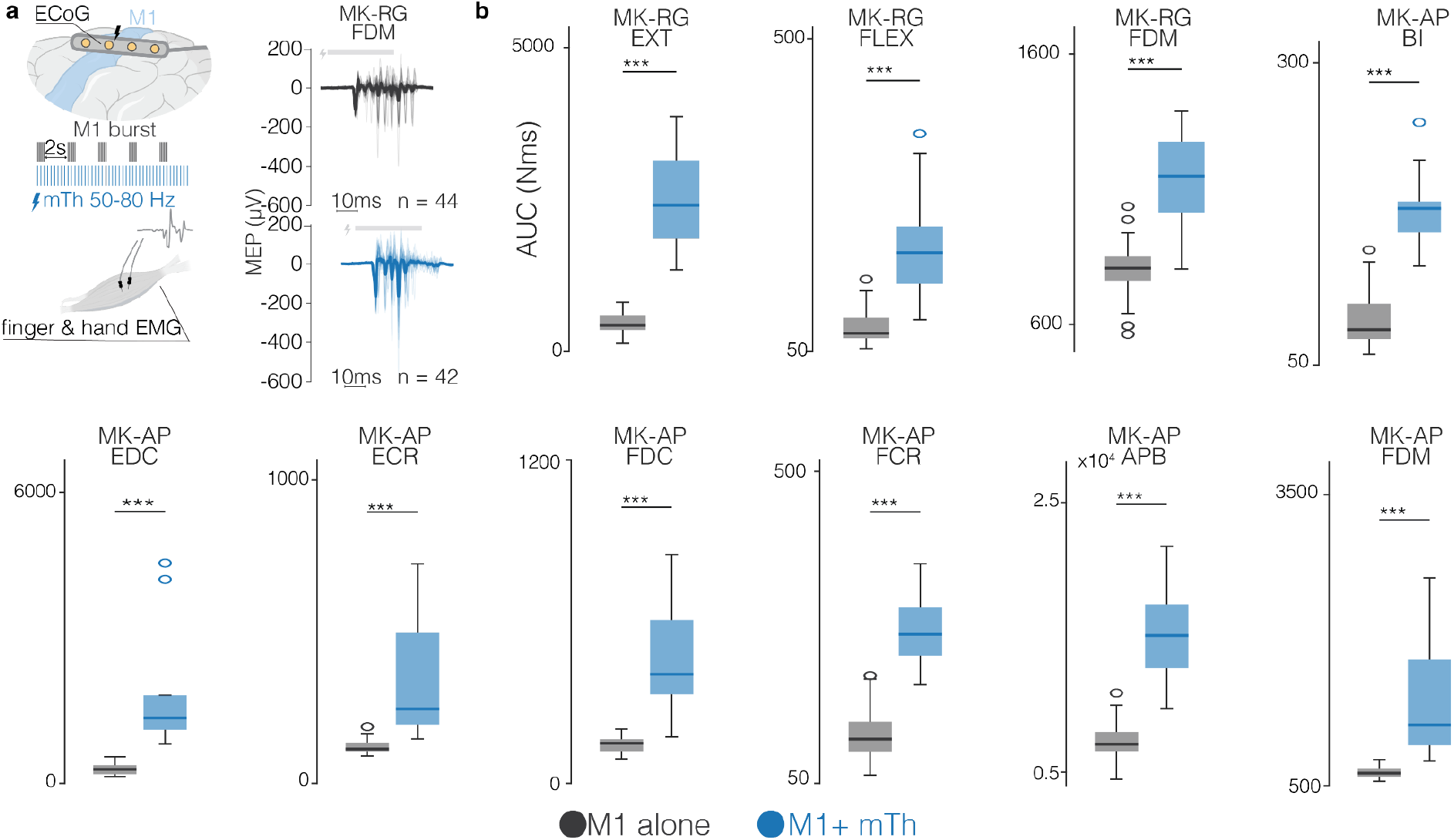
Potentiation of MEPs elicited by M1 stimulation. (**a**) <u>Left:</u> schematic of experimental setup. Burst of M1 stimulation were delivered through a subdural strip placed over M1. EMGs were recorded from upper limb muscles with and without low-frequency mTh stimulation. <u>Right</u>: example of EMG recordings following M1 burst stimulation, without (black) and with (blue) stimulations of the mTh. (**b**) for all animals and all muscles, box-plot of the area under the curve of EMG envelopes following M1 burst stimulation, without (black) and with (blue) stimulations of the mTh. We report only muscles that showed responses to stimulation. For all boxplots, the whiskers extend to the maximum spread not considering outliers, central, top, and bottom lines represent median, 25th, and 75th percentile, respectively.

**Extended Data Figure 3:**
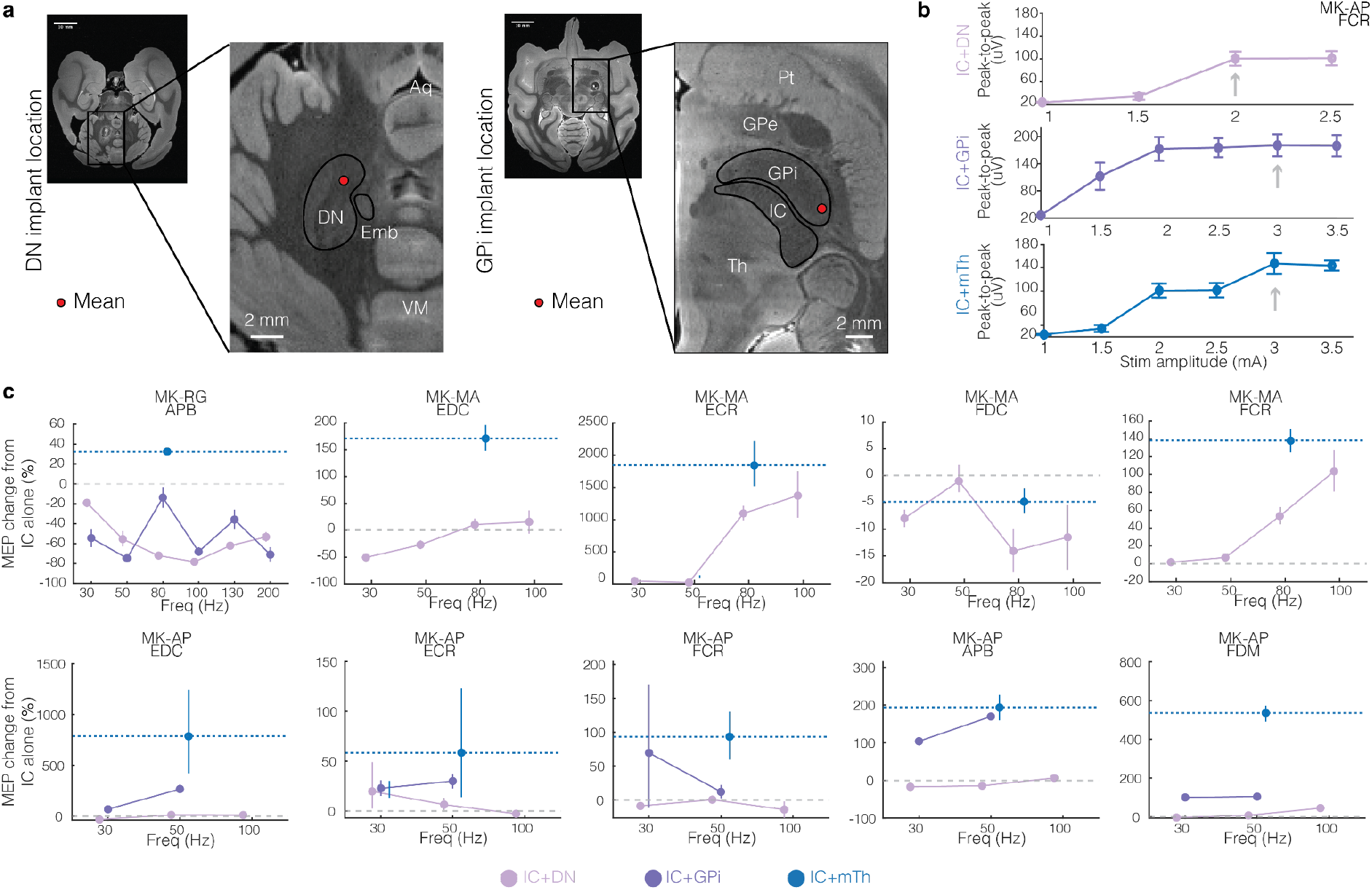
potentiation of MEPs with mTh, DN and GPi. (**a**) MRI showing reconstructed DN (left) and GPi (right) stimulation location, shown as average across animals. Black line shows the anatomical border of the nucleus. (**b**) Example of representative recruitment curve obtained by stimulating the IC with the mTh, DN or GPi at multiple amplitudes, until a MEP amplitude plateau is reached. Gray arrows indicate the amplitude used for MEP potentiation experiments. (**c**) For all animals and muscles, percentage change in MEP amplitude relative to IC alone as a function of stimulation frequency for stimulation of the DN (pink) and the GPi (purple) and compared to the maximum potentiation achieved with low-frequency mTh stimulation (dashed blue line). We report only muscles that showed responses to stimulation Data shown as mean ± 95% CI.

**Extended Data Figure 4:**
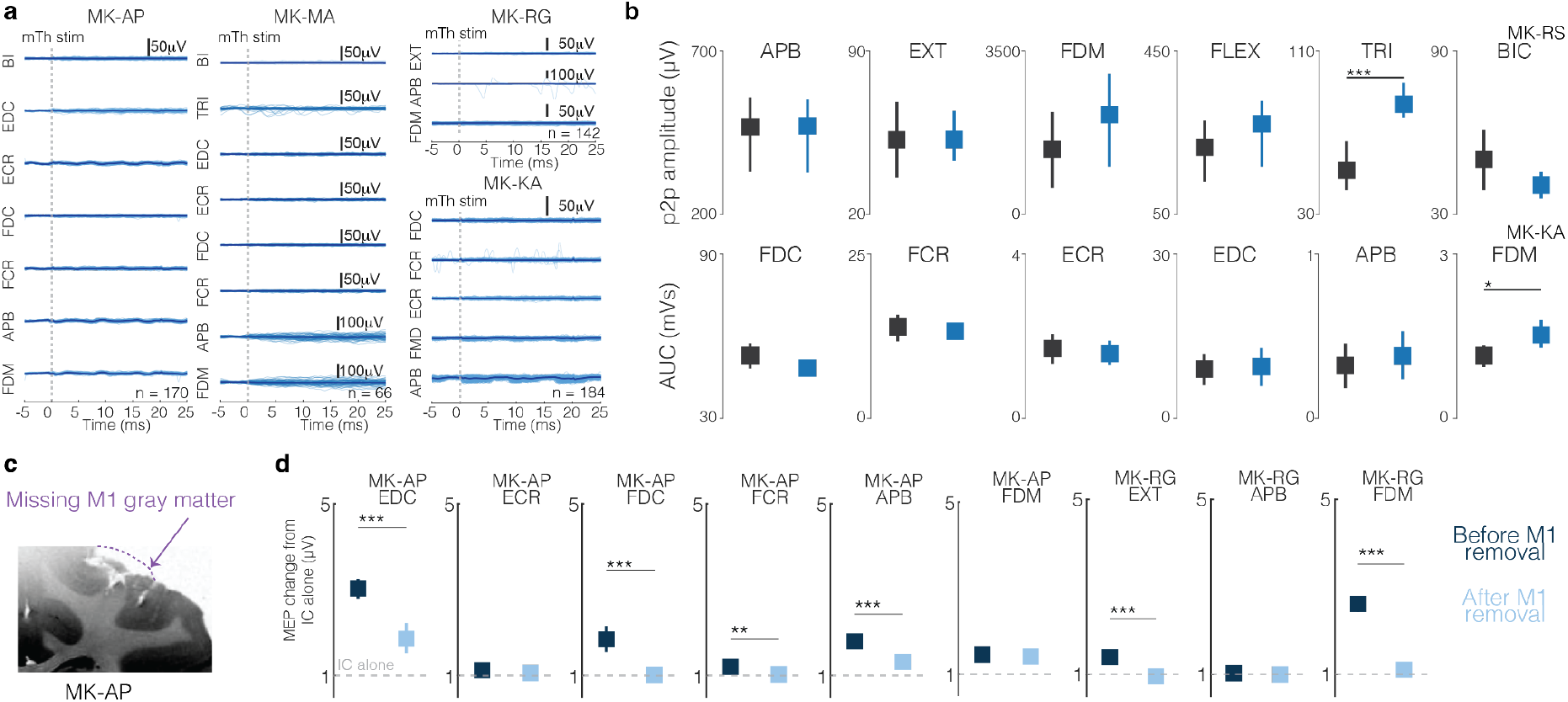
Electrophysiology shows that potentiation of MEPs occurs via increased thalamocortical synaptic input to M1 corticospinal neurons. (**a**) Potentiation of M1-induced MEPs. <u>Left</u>: schematic of experimental setup. Burst of M1 stimulation were delivered through a subdural strip placed over M1. EMGs were recorded from upper limb muscles with and without low-frequency mTh stimulation. <u>Middle</u>: example of EMG recordings following M1 burst stimulation, without (black) and with (blue) stimulations of the mTh. <u>Right</u>: for all animals and all muscles, box-plot of the area under the curve of EMG envelopes following M1 burst stimulation, without (black) and with (blue) stimulations of the mTh. For all boxplots, the whiskers extend to the maximum spread not considering outliers, central, top, and bottom lines represent median, 25th, and 75th percentile, respectively. (**b**) For all animals and all muscles, EMG traces with mTh stimulation alone at 50 Hz. (**c**) For all animals and all muscles, peak-to-peak amplitude or AUC of H-reflex elicited by dorsal root stimulation without (black) and with (blue) mTh stimulation. (**d**) Post-mortem MRI showing dissected M1. (**e**) Peak-to-peak amplitude change with mTh (normalized to IC stimulation alone) before and after surgical dissection of M1, for all animals and all muscles. Data shown as mean ± 95% CI. We report only muscles that showed responses to stimulation. For all panels, statistical significance was assessed with two-tail bootstrapping with Bonferroni correction: p<0.05 (*), p<0.01 (**), p<0.001(***).

**Extended Data Figure 5:**
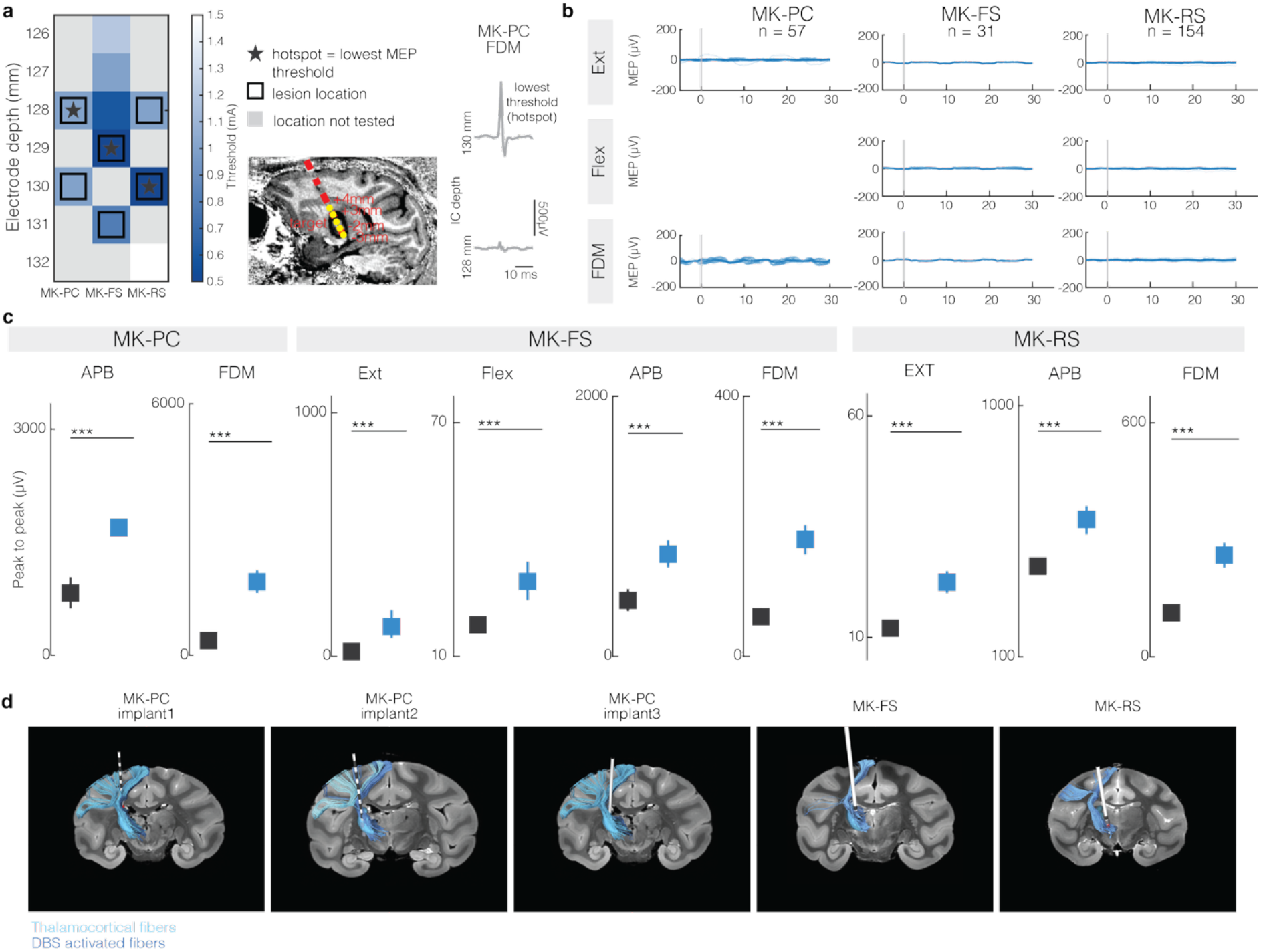
Surgical procedure to induce a CST lesion and to implant a chronic DBS lead. (**a**) Heatmap of MEPs at different depths along the IC trajectory, highlighting the hotspot (location with lowest motor threshold to elicit a MEP) where we induced the lesion. On the left, example of MEPs at two representative depths. (**b**) For all animals and all muscles, EMG traces with mTh DBS alone at 50 Hz. (**c**) Peak-to-peak MEP amplitude for all muscles and all animals to confirm implant location. Data shown as mean ± 95% CI. (**d**) For all monkeys and all implants, 3D reconstruction of DBS implant within the mTh (blue nucleus). Thalamocortical (mTh to M1) and DBS activated (activation volume to M1) fibers reconstructed from HDFT are shown. For all panels, statistical significance was assessed with two-tail bootstrapping with Bonferroni correction: p<0.05 (*), p<0.01 (**), p<0.001(***).

**Extended Data Figure 6:**
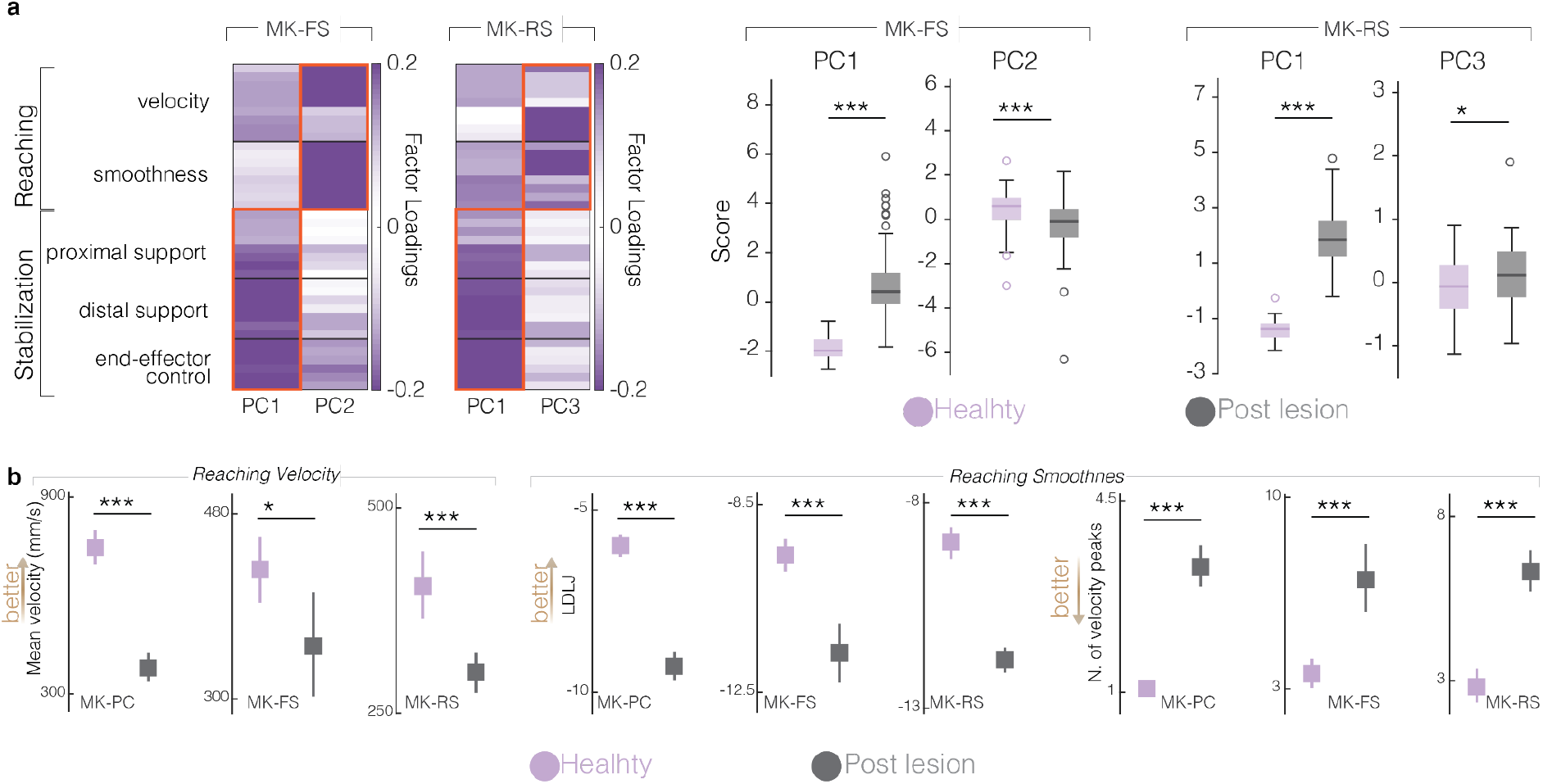
CST lesion effect on *touchscreen* and r*each, grasp & pull tas*k. (**a**) Loadings of features grouped according to their role in motor control for MK-FS and MK-RS. On the right, box-plot showing the PC scores before and after the lesion, for the first two PCs separating healthy and post-lesion reaches for MK-FS and MK-RS. For all boxplots, the whiskers extend to the maximum spread not considering outliers, central, top, and bottom lines represent median, 25th, and 75th percentile, respectively. (**b**) Kinematic features for the reaching component of the *reach, grasp & pull* task. All features are calculated from the wrist marker. For all panels, statistical significance was assessed with two-tail bootstrapping with Bonferroni correction: p<0.05 (*), p<0.01 (**), p<0.001(***).

**Extended Data Figure 7:**
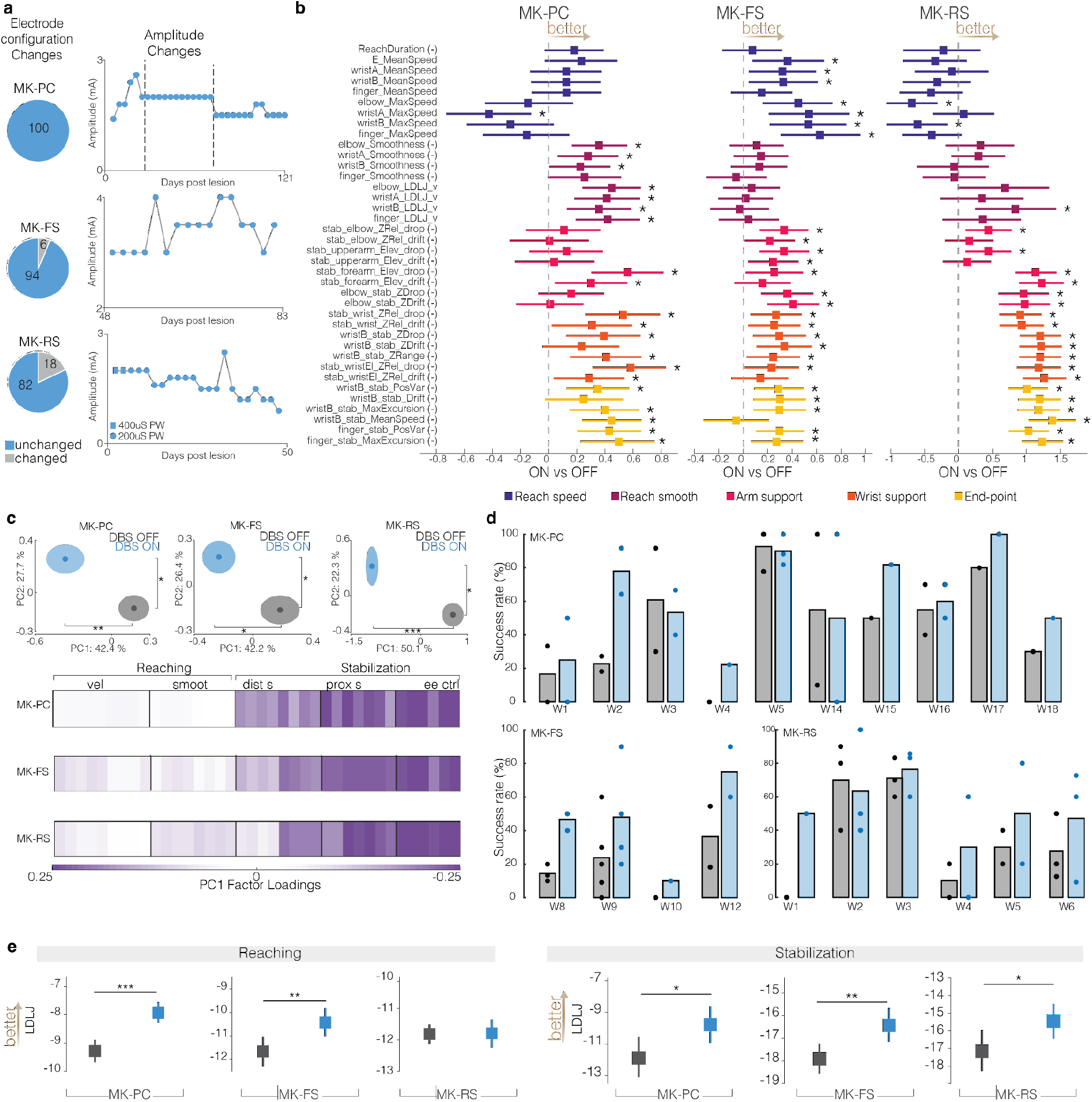
DBS benefit on the touchscreen and reach, grasp & pull task. (**a**) pie charts of sessions with changes in electrode configuration (location of stimulation), and scatter plots showing the changes of mTh stimulation amplitudes over time. (**b**) Complete list of the 38 kinematic features calculated for the touchscreen task. Data are shown as mean ± 95%CI of the DBS ON movements, normalized for the median DBS OFF. Sign indicates features are changed so that positive values represent a better outcome compared to DBS OFF. Asterisks indicate statistical significance, see methods paragraph. (**c**) PCA space showing separation of DBS ON and DBS OFF movements for all animals. On the bottom, the loading factors of the first PC highlight the critical effect of DBS on the stabilization phase of movement. (**d**) Bar plot of task success rate of the *reach, grasp & pull* task for all animals. (**e**) Representative features of the reaching and stabilization component of the *reach, grasp & pull* task. For all panels, statistical significance was assessed with two-tail bootstrapping with Bonferroni correction: p<0.05 (*), p<0.01 (**), p<0.001(***).

**Extended Data Figure 8:**
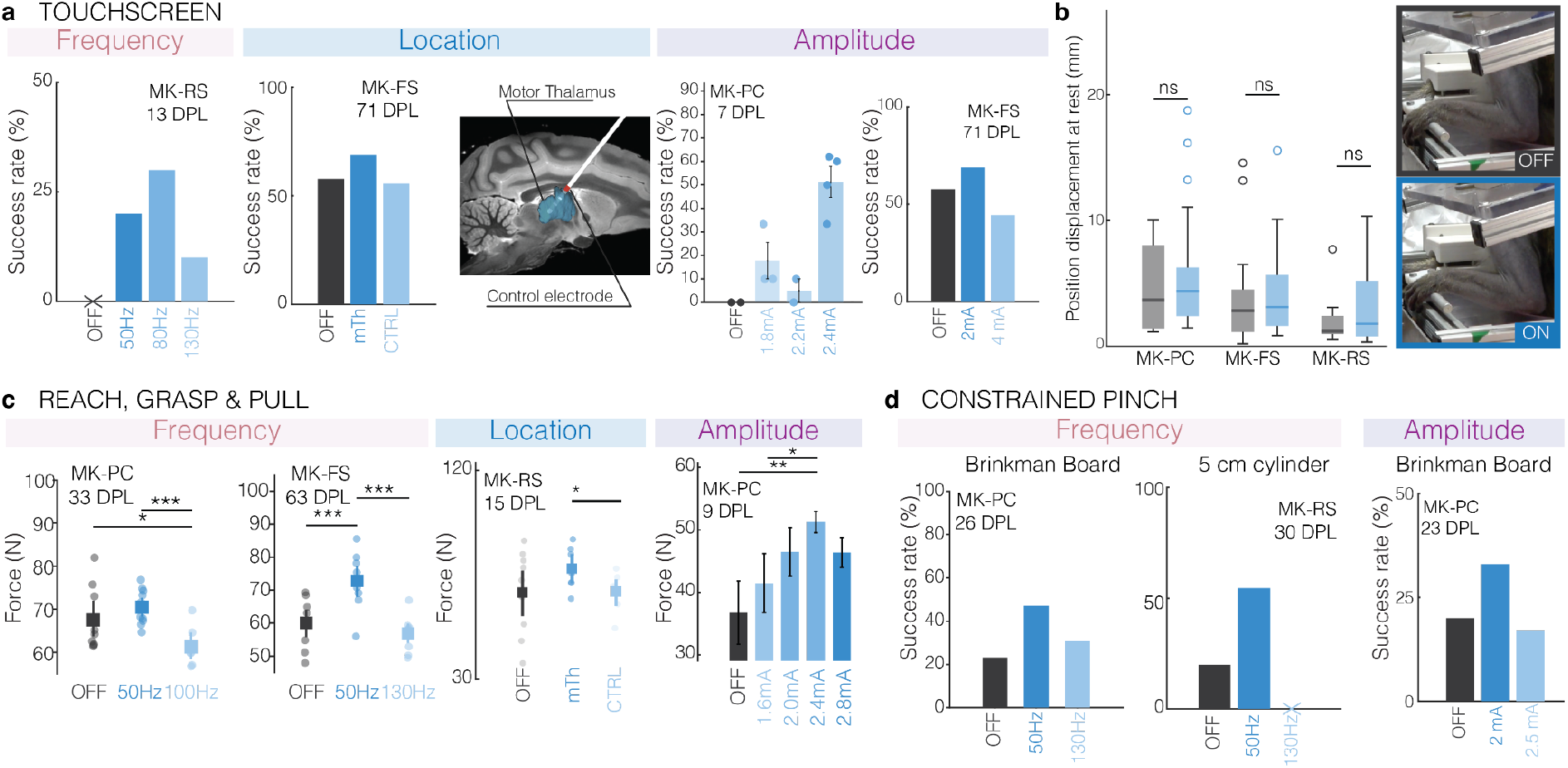
performance under control conditions with varying stimulation frequencies and location. (**a**) Effect on task success rate for multiple tested frequencies (left) and stimulation location (center) and amplitude (right). For the location, the snapshot shows in red the contact used for stimulation, outside of the mTh. (**b**) <u>Left:</u> Boxplot of 3D position displacement of the wrist marker during the resting period in the touchscreen task. For all boxplots, the whiskers extend to the maximum spread not considering outliers, central, top, and bottom lines represent median, 25th, and 75th percentile, respectively. <u>Right:</u> example of snapshots showing the resting position of MK-PC with DBS OFF (top) and DBS ON (bottom). No difference was found. (**c**) as in a, for the reach, grasp & pull task. (**d**) as in a and c, for the *constrained pinch* task. For all panels, statistical significance was assessed with two-tail bootstrapping with Bonferroni correction: p<0.05 (*), p<0.01 (**), p<0.001(***).

## SUPPLEMENTARY TABLE

**Supplementary Table 1.** Experimental procedures on each monkey. X= performed. nX = performed n times. -= not performed. * MK-RS dorsal root stimulation experiments were performed during an anesthetized procedure performed to implant a spinal array at day 50 post-lesion, as part of a different project.

|  | **Preoperative imaging** |  | **Behavioral testing** |  |  | **Terminal procedure** |  |  |  |  |  |
|---|---|---|---|---|---|---|---|---|---|---|---|
|  | **MRI** | **CT scan and fiducial screws** | **Behavioral training** | **Chronic mTh-DBS implant** | **CST lesion** | **mTh probe** | **GPi probe** | **DN probe** | **dorsar root stimulation** | **IC probe** | **ECoG** |
| **MK-PC** | X | X | X | 3X | X | X | - | - | - | X | - |
| **MK-RS** | X | X | X | X | X | X | - | - | X* | - | - |
| **MK-FS** | X | X | X | 2X | X | X | - | - | - | X | - |
| **MK-AP** | X | X | - | - | - | X | X | X | - | X | X |
| **MK-RG** | X | X | - | - | - | X | X | X | - | X | X |
| **MK-MA** | X | X | - | - | - | X | - | X | - | X | - |
| **MK-KA** | X | X | - | - | - | X | - | - | X | X | - |

**Supplementary Table 2.**
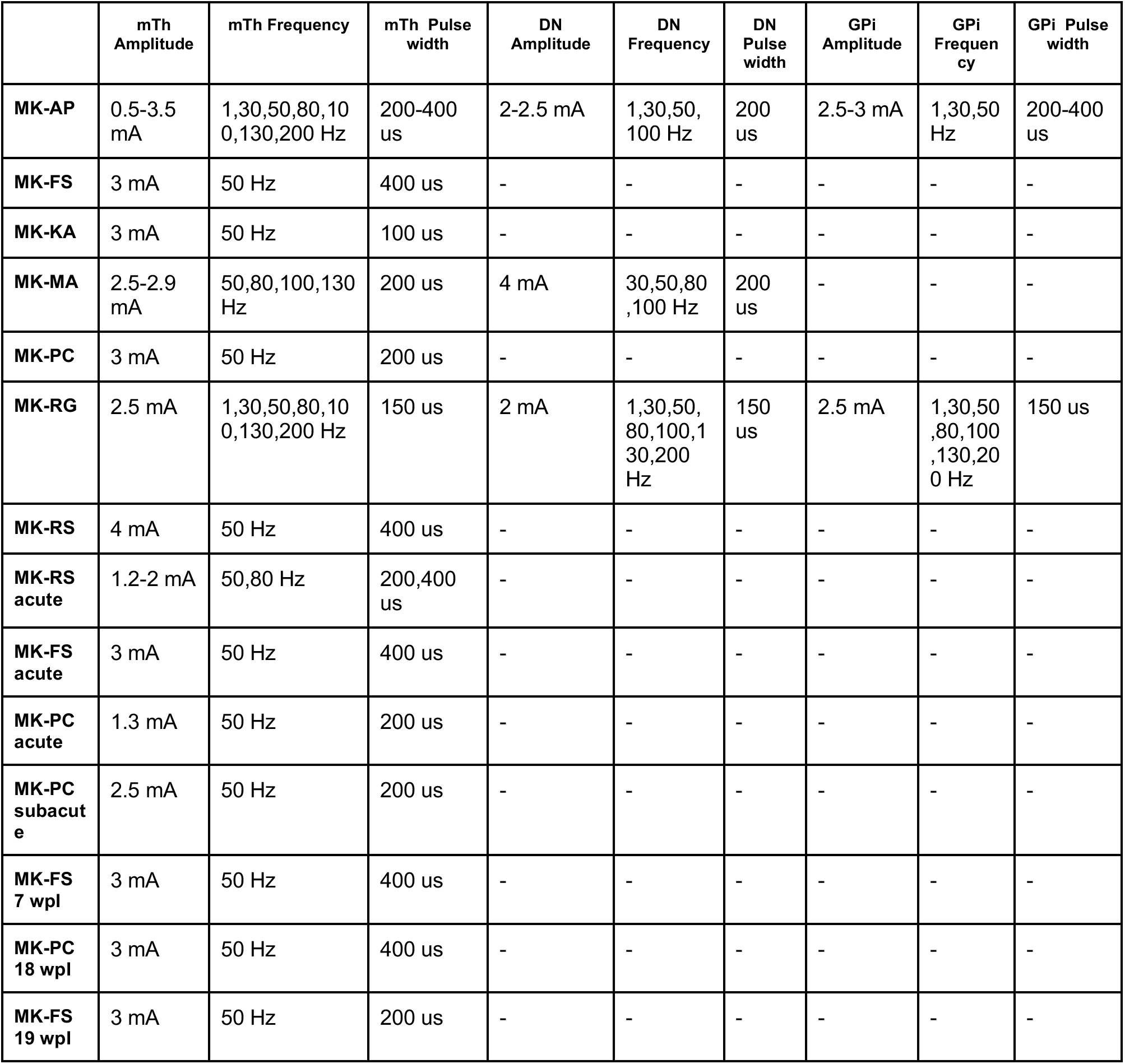
Stimulation parameters in all anesthetized experiments. These parameters include all experiments shown in Figures 1–2–3 and Extended Data Figures 1-2-3-4.

|  | **mTh Amplitude** | **mTh Frequency** | **mTh Pulse width** | **DN Amplitude** | **DN Frequency** | **DN Pulse width** | **GPi Amplitude** | **GPi Frequency** | **GPi Pulse width** |
|---|---|---|---|---|---|---|---|---|---|
| **MK-AP** | 0.5-3.5 mA | 1,30,50,80,100,130,200 Hz | 200-400 us | 2-2.5 mA | 1,30,50,100 Hz | 200 us | 2.5-3 mA | 1,30,50 Hz | 200-400 us |
| **MK-FS** | 3 mA | 50 Hz | 400 us | - | - | - | - | - | - |
| **MK-KA** | 3 mA | 50 Hz | 100 us | - | - | - | - | - | - |
| **MK-MA** | 2.5-2.9 mA | 50,80,100,130 Hz | 200 us | 4 mA | 30,50,80,100 Hz | 200 us | - | - | - |
| **MK-PC** | 3 mA | 50 Hz | 200 us | - | - | - | - | - | - |
| **MK-RG** | 2.5 mA | 1,30,50,80,100,130,200 Hz | 150 us | 2 mA | 1,30,50,80,100,130,200 Hz | 150 us | 2.5 mA | 1,30,50,80,100,130,200 Hz | 150 us |
| **MK-RS** | 4 mA | 50 Hz | 400 us | - | - | - | - | - | - |
| **MK-RS acute** | 1.2-2 mA | 50,80 Hz | 200,400 us | - | - | - | - | - | - |
| **MK-FS acute** | 3 mA | 50 Hz | 400 us | - | - | - | - | - | - |
| **MK-PC acute** | 1.3 mA | 50 Hz | 200 us | - | - | - | - | - | - |
| **MK-PC subacute** | 2.5 mA | 50 Hz | 200 us | - | - | - | - | - | - |
| **MK-FS 7 wpl** | 3 mA | 50 Hz | 400 us | - | - | - | - | - | - |
| **MK-PC 18 wpl** | 3 mA | 50 Hz | 400 us | - | - | - | - | - | - |
| **MK-FS 19 wpl** | 3 mA | 50 Hz | 200 us | - | - | - | - | - | - |

**Supplementary Table 3.** Kinematic features calculated for PCA analysis of touchscreen task. . Notation. t1, t2, t3: manually placed events (movement onset, arrival at the screen, feedback); fs: sampling rate. Reaching window: t ∈ [t1, t2]; stabilization window: t ∈ [t2, t3]. PM(t) = [x, y, z] position of marker M (shoulder, elbow, wristA, wristB, finger); X = forward, Y = lateral, Z = vertical (up positive). sM(t) = ∥dPM/dt∥ is the marker speed; ∥·∥ is the Euclidean norm. TR = reach duration; speak = max sM(t) in the reaching window. Δzse, Δzew (and Δx, Δy) are the coordinate differences elbow − shoulder and wristB − elbow, respectively. Because targets differ across trials, angles and heights are expressed only as changes relative to the first stabilization frame (t2). Distances in marker units (mm); angles in degrees.

| Group | Variable | Description | Formula |
|---|---|---|---|
| Reaching – velocity | ReachDuration | Movement time of the reach, from movement onset to arrival at the screen (higher = worse). | T<sub>R</sub> = (t<sub>2</sub> − t<sub>1</sub>) / f<sub>s</sub> |
|  | elbow_MeanSpeed | Mean speed of the elbow marker during the reach, computed as net displacement divided by movement time (higher = better). | ‖P<sub>elbow</sub>(t<sub>2</sub>) − P<sub>elbow</sub>(t<sub>1</sub>)‖ / T<sub>R</sub> |
|  | wristA_MeanSpeed | Mean speed of the wristA marker during the reach, computed as net displacement divided by movement time (higher = better). | ‖P<sub>wristA</sub>(t<sub>2</sub>) − P<sub>wristA</sub>(t<sub>1</sub>)‖ / T<sub>R</sub> |
|  | wristB_MeanSpeed | Mean speed of the wristB marker during the reach, computed as net displacement divided by movement time (higher = better). | ‖P<sub>wristB</sub>(t<sub>2</sub>) − P<sub>wristB</sub>(t<sub>1</sub>)‖ / T<sub>R</sub> |
|  | finger_MeanSpeed | Mean speed of the finger marker during the reach, computed as net displacement divided by movement time (higher = better). | ‖P<sub>finger</sub>(t<sub>2</sub>) − P<sub>finger</sub>(t<sub>1</sub>)‖ / T<sub>R</sub> |
|  | elbow_MaxSpeed | Peak speed of the elbow marker during the reach (higher = better). | max<sub>t∈[t<sub>1</sub>,t<sub>2</sub>]</sub> s<sub>elbow</sub>(t) |
|  | wristA_MaxSpeed | Peak speed of the wristA marker during the reach (higher = better). | max<sub>t∈[t<sub>1</sub>,t<sub>2</sub>]</sub> s<sub>wristA</sub>(t) |
|  | wristB_MaxSpeed | Peak speed of the wristB marker during the reach (higher = better). | max<sub>t∈[t<sub>1</sub>,t<sub>2</sub>]</sub> s<sub>wristB</sub>(t) |
|  | finger_MaxSpeed | Peak speed of the finger marker during the reach (higher = better). | max<sub>t∈[t<sub>1</sub>,t<sub>2</sub>]</sub> s<sub>finger</sub>(t) |
| Reaching – smoothness | elbow_Smoothness | Number of peaks in the elbow speed profile during the reach; more peaks indicate a more fragmented movement (higher = worse). | N<sub>peaks</sub>[ s<sub>elbow</sub>(t) ], t ∈ [t<sub>1</sub>, t<sub>2</sub>] |
|  | wristA_Smoothness | Number of peaks in the wristA speed profile during the reach; more peaks indicate a more fragmented movement (higher = worse). | N<sub>peaks</sub>[ s<sub>wristA</sub>(t) ], t ∈ [t<sub>1</sub>, t<sub>2</sub>] |
|  | wristB_Smoothness | Number of peaks in the wristB speed profile during the reach; more peaks indicate a more fragmented movement (higher = worse). | N<sub>peaks</sub>[ s<sub>wristB</sub>(t) ], t ∈ [t<sub>1</sub>, t<sub>2</sub>] |
|  | finger_Smoothness | Number of peaks in the finger speed profile during the reach; more peaks indicate a more fragmented movement (higher = worse). | N<sub>peaks</sub>[ s<sub>finger</sub>(t) ], t ∈ [t<sub>1</sub>, t<sub>2</sub>] |
|  | elbow_LDLJ_v | Log dimensionless jerk of the elbow speed profile during the reach (higher = smoother). | −ln( T<sub>R</sub><sup>3</sup> / s<sub>peak</sub><sup>2</sup> · ∫ (d²s<sub>elbow</sub>/dt²)² dt ), t ∈ [t<sub>1</sub>, t<sub>2</sub>] |
|  | wristA_LDLJ_v | Log dimensionless jerk of the wristA speed profile during the reach (higher = smoother). | −ln( T<sub>R</sub><sup>3</sup> / s<sub>peak</sub><sup>2</sup> · ∫ (d²s<sub>wristA</sub>/dt²)² dt ), t ∈ [t<sub>1</sub>, t<sub>2</sub>] |
|  | wristB_LDLJ_v | Log dimensionless jerk of the wristB speed profile during the reach (higher = smoother). | −ln( T<sub>R</sub><sup>3</sup> / s<sub>peak</sub><sup>2</sup> · ∫ (d²s<sub>wristB</sub>/dt²)² dt ), t ∈ [t<sub>1</sub>, t<sub>2</sub>] |
|  | finger_LDLJ_v | Log dimensionless jerk of the finger speed profile during the reach (higher = smoother). | −ln( T<sub>R</sub><sup>3</sup> / s<sub>peak</sub><sup>2</sup> · ∫ (d²s<sub>finger</sub>/dt²)² dt ), t ∈ [t<sub>1</sub>, t<sub>2</sub>] |
| Stabilization – proximal support | stab_elbow_ZRel_drop | Maximal drop of the elbow height relative to the shoulder, with respect to the first stabilization frame (higher = worse). | Δ<sub>es</sub>(t<sub>2</sub>) − min<sub>t</sub> Δ<sub>es</sub>(t), Δ<sub>es</sub> = z<sub>elbow</sub> − z<sub>shoulder</sub> |
|  | stab_elbow_ZRel_drift | Net drop of the elbow height relative to the shoulder between the first and last stabilization frame (higher = worse). | Δ<sub>es</sub>(t<sub>2</sub>) − Δ<sub>es</sub>(t<sub>3</sub>) |
|  | stab_upperarm_Elev_drop | Maximal loss of upper-arm elevation angle (shoulder→elbow) within the stabilization window, relative to the first frame; angular change only, deg (higher = worse). | θ<sub>UA</sub>(t<sub>2</sub>) − min<sub>t</sub> θ<sub>UA</sub>(t), θ<sub>UA</sub> = atan2( Δz<sub>se</sub>, √(Δx<sub>se</sub>² + Δy<sub>se</sub>²) ) |
|  | stab_upperarm_Elev_drift | Net loss of upper-arm elevation angle between the first and last stabilization frame, deg (higher = worse). | θ<sub>UA</sub>(t<sub>2</sub>) − θ<sub>UA</sub>(t<sub>3</sub>) |
|  | stab_forearm_Elev_drop | Maximal loss of forearm elevation angle (elbow→wrist B) within the stabilization window, relative to the first frame, deg (higher = worse). | θ<sub>FA</sub>(t<sub>2</sub>) − min<sub>t</sub> θ<sub>FA</sub>(t), θ<sub>FA</sub> = atan2( Δz<sub>ew</sub>, √(Δx<sub>ew</sub>² + Δy<sub>ew</sub>²) ) |
|  | stab_forearm_Elev_drift | Net loss of forearm elevation angle between the first and last stabilization frame, deg (higher = worse). | θ<sub>FA</sub>(t<sub>2</sub>) − θ<sub>FA</sub>(t<sub>3</sub>) |
|  | elbow_stab_ZDrop | Maximal vertical drop of the elbow marker during stabilization (absolute Z; includes the shoulder/trunk contribution) (higher = worse). | z<sub>elbow</sub>(t<sub>2</sub>) − min<sub>t</sub> z<sub>elbow</sub>(t) |
|  | elbow_stab_ZDrift | Net vertical drop of the elbow marker between the first and last stabilization frame (absolute Z) (higher = worse). | z<sub>elbow</sub>(t<sub>2</sub>) − z<sub>elbow</sub>(t<sub>3</sub>) |
| Stabilization – distal support | stab_wrist_ZRel_drop | Maximal drop of wrist B height relative to the shoulder, with respect to the first stabilization frame (higher = worse). | Δ<sub>ws</sub>(t<sub>2</sub>) − min<sub>t</sub> Δ<sub>ws</sub>(t), Δ<sub>ws</sub> = z<sub>wristB</sub> − z<sub>shoulder</sub> |
|  | stab_wrist_ZRel_drift | Net drop of wrist B height relative to the shoulder between the first and last stabilization frame (higher = worse). | Δ<sub>ws</sub>(t<sub>2</sub>) − Δ<sub>ws</sub>(t<sub>3</sub>) |
|  | wristB_stab_ZDrop | Maximal vertical drop of wrist B during stabilization (absolute Z; includes shoulder and elbow contributions) (higher = worse). | z<sub>wristB</sub>(t<sub>2</sub>) − min<sub>t</sub> z<sub>wristB</sub>(t) |
|  | wristB_stab_ZDrift | Net vertical drop of wrist B between the first and last stabilization frame (absolute Z) (higher = worse). | z<sub>wristB</sub>(t<sub>2</sub>) − z<sub>wristB</sub>(t<sub>3</sub>) |
|  | wristB_stab_ZRange | Total vertical excursion of wrist B during stabilization (absolute Z) (higher = worse). | max<sub>t</sub> z<sub>wristB</sub>(t) − min<sub>t</sub> z<sub>wristB</sub>(t) |
|  | stab_wristEl_ZRel_drop | Maximal drop of wrist B height relative to the elbow, isolating wrist-only support (higher = worse). | Δ<sub>we</sub>(t<sub>2</sub>) − min<sub>t</sub> Δ<sub>we</sub>(t), Δ<sub>we</sub> = z<sub>wristB</sub> − z<sub>elbow</sub> |
|  | stab_wristEl_ZRel_drift | Net drop of wrist B height relative to the elbow between the first and last stabilization frame (higher = worse). | Δ<sub>we</sub>(t<sub>2</sub>) − Δ<sub>we</sub>(t<sub>3</sub>) |
| Stabilization – end-effector control | wristB_stab_PosVar | Positional variance of wrist B (3D) during stabilization (higher = worse). | ( σ²<sub>x</sub> + σ²<sub>y</sub> + σ²<sub>z</sub>) / 3, of P<sub>wristB</sub>(t), t ∈ [t<sub>2</sub>, t<sub>3</sub>] |
|  | wristB_stab_Drift | Net 3D displacement of wrist B between the first and last stabilization frame (higher = worse). | ‖P<sub>wristB</sub>(t<sub>3</sub>) − P<sub>wristB</sub>(t<sub>2</sub>)‖ |
|  | wristB_stab_MaxExcursion | Maximal 3D distance of wrist B from its arrival position during stabilization (higher = worse). | max<sub>t</sub> ‖P<sub>wristB</sub>(t) − P<sub>wristB</sub>(t<sub>2</sub>)‖ |
|  | wristB_stab_MeanSpeed | Mean residual speed of wrist B during stabilization (higher = worse). | mean<sub>t</sub> s<sub>wristB</sub>(t), t ∈ [t<sub>2</sub>, t<sub>3</sub>] |
|  | finger_stab_PosVar | Positional variance of the finger (3D) during stabilization (higher = worse). | ( σ²<sub>x</sub> + σ²<sub>y</sub> + σ²<sub>z</sub>) / 3, of P<sub>finger</sub>(t), t ∈ [t<sub>2</sub>, t<sub>3</sub>] |
|  | finger_stab_MaxExcursion | Maximal 3D distance of the finger from its arrival position during stabilization (higher = worse). | max<sub>t</sub> ‖P<sub>finger</sub>(t) − P<sub>finger</sub>(t<sub>2</sub>)‖ |

